# Small RNA-mediated genome silencing in *Caenorhabditis elegans* promotes telomere stability in the absence of telomerase

**DOI:** 10.1101/292722

**Authors:** Lu Lu, Ambika Bhattarai, Charlie Longtine, Stephen Frenk, Madelyn Garrison, Julie Seohyun Lee, Evan H. Lister-Shimauchi, Shawn Ahmed

## Abstract

We previously found that small RNAs with perfect homology to (TTAGGC)_n_ telomere repeats are very rare. Here we use long read sequencing to reveal that most N2 wild type *C. elegans* strains have few telomere mutations at 5’ ends of telomeres, where the telomeric lncRNA TERRA is transcribed. Therefore, telomere mutations cannot explain the absence of perfect small RNAs to (TTAGGC)_n_ telomere repeats. We show that the heterochromatin mark H3K9me2 is enriched at *C. elegans* sub-telomeres and that the HP1 protein HPL-2 that binds H3K9me2 promotes telomere stability in the absence of telomerase. Small RNA silencing can promote H3K9 methylation, and telomere stability in the absence of telomerase requires germ granule proteins MUT-7 and MUT-14 that promote small RNA biogenesis and also WAGO-1 and HRDE-1 Argonaute proteins that bind subtelomeric small RNAs. Loss of telomerase induced biogenesis of small RNAs from the subtelomeric 5’ end of the telomeric lncRNA TERRA but not from telomere repeats of TERRA. Loss of both telomerase and small RNA-mediated genome silencing induced TERRA expression, telomere damage and accelerated sterility. We propose that small RNA-mediated heterochromatin formation and telomerase function redundantly to repress a form of telomeric DNA damage that is coupled to TERRA expression and subtelomeric small RNA biogenesis. However, telomere repeats of TERRA strongly resisted RNA-dependent RNA polymerase activity in telomerase mutants, revealing an exceptional quality of telomere repeat RNA.

**Article summary:** Loss of telomerase and either small RNA biogenesis factors or heterochromatin proteins resulted in rapid onset of telomere fusions. We identified Argonaute proteins that bind to subtelomeric small RNAs and promote telomere stability in the absence of telomerase. Telomeres are transcribed to create the non-coding RNA TERRA that is a composite of subtelomere sequence and telomere repeats. Although subtelomeric small RNAs were upregulated in the absence of telomerase, the telomere sequence of TERRA was completely resistant to small RNA biogenesis. Small RNAs and TERRA may interact to promote repair of damaged telomeres that cannot be healed by telomerase.

## Introduction

Telomeres are repetitive nucleoprotein structures that protect the ends of linear eukaryotic chromosomes from degradation and recognition as double-stranded DNA breaks (Maciejowski and de Lange 2017; Shay and Wright 2019). Deficiency for telomerase reverse transcriptase in most human somatic cells results in progressive telomere erosion over cell divisions, which eventually triggers replicative senescence (Micco et al. 2021). Telomere shortening acts as a barrier to tumorigenesis, but may also contribute to organismal aging through impaired stem cell function, regeneration, and organ maintenance (Donate and Blasco 2011; López-Otín et al. 2023). Regulating the rate of telomere erosion in telomerase-negative cells is therefore crucial for maintaining the balance between tumor suppression and age-related tissue decline. Tumor cells escape telomere-triggered senescence by stabilizing telomere length, either through reactivation of telomerase or through activation of the telomerase-independent telomere maintenance pathway termed Alternative Lengthening of Telomeres (ALT) (Pickett and Reddel 2015).

Telomeres have features characteristic of heterochromatin in that they associate with the heterochromatin protein HP1 and contain transcriptionally repressive histone marks such as methylation of H3K9 and H4K20 (Benetti et al. 2007; McMurchy et al. 2017; Bettin et al. 2025; van der Lugt and Jacobs 2026). Telomeres are transcribed to create the G-rich long noncoding RNA (lncRNA) termed TERRA, a feature widely conserved among eukaryotes, and lower levels of the C-rich antisense lncRNA ARRET have also been reported (Azzalin et al. 2007; Abreu et al. 2025). TERRA is transcribed from the subtelomere into the telomere by RNA polymerase II and is upregulated at shortened and damaged telomeres, where it may recruit telomerase and chromatin modifiers to promote telomere elongation (Cusanelli et al. 2013; Azzalin and Lingner 2015; Rivosecchi et al. 2024). TERRA expression in mouse cells has been linked to H3K9me3 and other heterochromatic histone marks present at mammalian telomeres (Montero et al. 2018). Telomeric small RNAs likely created from TERRA have been observed in *Tetrahymena* and mouse stem cells (Cao et al. 2009; Couvillion et al. 2009), and they are associated with the presence of subtelomeric DNA methylation in *Arabidopsis* (Vrbsky et al. 2010).

Sites of DNA damage in mammals and *Arabidopsis* have been reported to create bidirectional transcripts that form dsRNA intermediates, and they could be processed by the RNase class III enzyme Dicer into small RNAs that may modify chromatin and promote homologous recombination (Francia et al. 2012; Wei et al. 2012; Sharma and Misteli 2013). Similarly, telomere uncapping in mammals induces expression of TERRA and ARRET, which are processed by Dicer into telomeric small RNAs that promote the DNA damage response at dysfunctional telomeres (Rossiello et al. 2017; Rossiello et al. 2022).

The *C. elegans* homologue of Dicer, DCR-1, can cleave exogenous stranded RNA to create primary sRNAs that trigger production of effector sRNAs that promote transcriptional silencing or mRNA degradation (Billi et al. 2018). In the *C. elegans* germline, mRNAs targeted by small RNA/Argonaute complexes are transported to Mutator bodies (Uebel et al. 2018), where RNA-dependent RNA polymerase (RdRP) uses target mRNA as a template to create 22 nucleotide sRNAs with a 5’ guanine (22G RNAs) (Pak and Fire 2007). These effector 22G RNAs are loaded onto Worm-specific Argonaute proteins (WAGOs) such as HRDE-1, a nuclear Argonaute protein that promotes heterochromatin deposition at loci targeted by small RNAs (Ashe et al. 2012; Buckley et al. 2012). Germline transcription is protected by a distinct category of 22G RNAs that interact with the anti-silencing Argonaute protein CSR-1 (Avgousti et al. 2012; Wedeles et al. 2013).

Here we describe an endogenous sRNA and heterochromatin pathway that functions redundantly with telomerase to repress TERRA and promote telomere stability. We show that telomere stability in the absence of telomerase depends on heterochromatin proteins SPR-5, RBR-2 and HPL-2 that promote H3K9 methylation, on HRDE-1 and WAGO-1 Argonaute proteins that interact with subtelomeric and telomeric 22G small RNAs, and on germ granule proteins MUT-7 and MUT-14 that promote biogenesis of these small RNAs.

## Materials and Methods

### Strains

Unless otherwise noted, all strains were maintained at 20°C on nematode growth medium plates seeded with *Escherichia coli* OP50. Strains used include Bristol N2 ancestral, *wago-1(tm1414)*, *trt-1(ok410), unc-29(e193), dpy-5(e61), dcr-1(mg375), hrde-1(tm1200), rde-1(ne219), rde-4(ne299), rrf-3(mg373), rrf-2(ok210), eri-1(mg366), mut-7(pk204), mut-7(pk720), mut-14(pk738), rbr-2(tm1231), spr-5(by101), xnp1(ok1823), xnp-1(tm678), hpl-1(tm1624), hpl-2(ok916), hpl-2(tm1479),* and *exo1(tm1842)*. A *trt-1 unc-29* double mutant, where *unc-29* is tightly linked to *trt-1*, was used to create strains used in this study. Double, triple, and quadruple mutants were created using standard marker mutations and verified based on phenotype or genotype. *mut-7* and *mut-14* mutants are known to accumulate transposon-induced mutations, so these strains were outcrossed several times before use.

### DAPI staining

One-day-old adults were washed twice in M9, soaked in 150 µL of 400ng/mL DAPI in ethanol solution for 30 min or until evaporated, rehydrated in 2 mL of M9 solution overnight at 4°C, and mounted in 8 µL VECTASHIELD mounting medium (Vector Laboratories). Chromosome counts were performed under 60x magnification and a 359nm excitation wavelength using a Nikon Eclipse E800 microscope.

### qRT-PCR

Non-starved mixed-stage worms were washed with M9 and RNA was extracted using 1 mL TRIZOL according to the manufacturer’s instructions. For RT-PCR analysis, total RNA was treated twice with DNase I and was reverse transcribed with primers for a reference gene *act-1* and a telomere-specific primer (GCCTAA)_3_ using SuperScript II reverse transcriptase (Invitrogen). For SYBR green reactions, subtelomere specific primers were designed within 200bp of the telomere-subtelomere junction. RT-PCR primer sequences are listed below.

IIIL forward: AAGCCTAAAAGCGCGAAATCC

IIIL reverse: TAGATTCCGTGCTTTCTCAGAC

VR forward: ATGGCGATAGAGGATACTGG

VR reverse: CCTAAGCAAATCCCCAGAAAGG

XL forward: AGAGTTCGACTTCGGGACAC

XL reverse: GCCTAATCTGTGCTCCAAAGCC

### RNA FISH

RNA FISH was performed as previously described (Sakaguchi et al. 2014) with several modifications. In brief, non-starved, mixed-stage animals were washed twice in M9 and three times in 1x DEPC-treated PBS. Worms were fixed in 1mL fixation buffer (1x DEPC-treated PBS with 3.7% formaldehyde) for 45 min rotating at room temperature then washed twice in 1mL 1x DEPC-treated PBS. Fixed animals were permeabilized overnight at 4°C in 70% ethanol in DEPC-treated H_2_O, then washed in 1 mL wash buffer [10% (vol/vol) formamide in 2x RNases-free SSC]. Aliquots of each sample were removed prior to hybridization and treated with RNaseA for 1h at 37°C to ensure probe specificity to RNA. Hybridization was performed at 30°C overnight in 100 µL hybridization buffer (10% dextran sulfate, 2x RNase-free SSC, 10% deionized formamide, 0.02% RNase-free BSA, 50 µg *E. coli* tRNA) with 1.25 µM probe. Cy5labelled (GCCTAA)_3_ probe was used to detect TERRA transcripts. Animals were washed twice with wash buffer, counterstained with 25 ng/mL DAPI for 30 min at room temperature, and mounted on glass slides in VECTASHIELD mounting medium (Vector Laboratories) before epifluorescence microscopy.

### Immunofluorescence

IF was carried out as previously described (Phillips et al., 2009) by fixing dissected gonads of 1 day old adults in 2% PFA for 10 minutes followed by a 1 minute postfixation in methanol. Rabbit anti-pS/TQ (Cell Signaling Technology antibody #2851) was used at a 1:500 dilution and mouse anti-RNA:DNA hybrid S9.6 (Cell Signaling Technology antibody #47099) at 1:500 dilution. Stained gonads were counterstained with DAPI and mounted with VECTASHIELD (Vector laboratories) before epifluorescence microscopy.

### RNAi

RNAi clone expressing dsRNA targeting *C. elegans* telomeres was constructed using NEB HiFi DNA assembly mix according to the manufacturer’s instructions. 200 bp of telomeric repeats were amplified from cTel55x by PCR with 20 bp overlap to SpeI digested L4440. Correct constructs were transformed into HT115 and RNAi was performed as previously described (McMurchy et al. 2017).

### Southern Blot

*C. elegans* genomic DNA was digested overnight with Hinf1 (New England Biolabs), separated on a 0.8% agarose gel at 1.5 V/cm, and transferred to a neutral nylon membrane (Hybond-N, GE Healthcare Life Sciences). A digoxigeninlabelled telomere probe was hybridized and detected as described (Ahmed and Hodgkin, 2000). Migration distance was measured using ImageJ.

### Mrt assay

Mortal Germline (Mrt) assay was conducted as previously described (Meier et al. 2006). Strains were passaged by transferring six L1 larval animals to fresh plates weekly, and the transgenerational sterility was observed. 20-40 plates per genotype were grown for Mrt assays, and Mantel-Cox Log Rank analysis was performed to compare transgenerational sterility between strains.

### Alt assay

Survival of telomerase mutants via the Alternative Lengthening of Telomeres (ALT) telomere maintenance pathway was assayed for *trt-1(ok410)* mutant strains by transferring a 1cm^2^ chunk of agar populated with ≈200–400 animals weekly. We previously observed that 10-20% of *trt-1* mutant strains survive for hundreds of generations. In contrast, 100% of sterility is observed within 20–30 generations when *trt-1* mutant strains were passaged via Mrt assays. A z-test was performed to compare the proportion of Allt survivors in chunked strains.

### RNA-seq

Animals were grown at 20°C on 100 mm NGM plates seeded with OP50 bacteria. Two biological replicates were prepared for each genotype. RNA was extracted using Trizol (Ambion) followed by isopropanol precipitation. RNA samples were treated with 20U RNA 5’ polyphosphatase (Lucigen, RP8092H) for 30 minutes at 37°C to remove 5’ triphosphate groups. Library preparation and sequencing was performed at the UNC School of Medicine High-Throughput Sequencing Facility (HTSF). RNA-seq libraries were prepared using the NEXTflex Small RNA Library Prep Kit for Illumina (Bio Scientific). Libraries were sequenced on an Illumina HiSeq 2500 to produce approximately twenty million 50 bp single-end reads per library.

### Analysis of high throughput sequencing data

Adapter trimming was performed as required using the bbduk.sh script from the bbmap suite3 and custom scripts (Bushnell, 2016). Reads were then filtered for 22G species or all species between 18 and 30 nucleotides in length using a custom python script. Reads were then converted to fasta format and mapped to the *C. elegans* genome (WS251) using bowtie (Langmead et al., 2009) with the following options: –M 1 –v 0 –best –strata. Uniquely mapping sRNAs were detected by replacing the –M 1 parameter with –m 1. To detect telomere-mapping sRNAs, reads were mapped to seven repeats of perfect telomere sequence (TTAGGC), allowing for zero mismatches. Unmapped reads were re-mapped to the telomere sequence, this time allowing for one mismatch. This step was repeated to detect reads that map to telomeres with two or three mismatches. Genes with differential sRNA abundance in *trt-1* were identified using DESeq2 (Love et al. 2014). To detect changes in the abundance of subtelomeric sRNAs in *trt-1* and *trt-1; dcr-1*, 22G reads mapping to the 2kb of sequence adjacent to the telomere were counted for each chromosome arm. Counts were normalized by dividing by the total number of mapped reads for each library. A pseudocount of 1 was added to each normalized value to avoid division by zero errors. Replicates were averaged and the log2-fold change between mutant and wildtype was calculated for each subtelomere. Analysis of sequencing data was performed using the R statistical computing environment (Team and Others 2017). Genome regions were visualized using the Gviz R package (Hahne and Ivanek 2016). To detect sRNAs targeting transposons in immunoprecipitation datasets, 22G reads were mapped to the *C. elegans* transposon consensus sequences downloaded from Repbase with bowtie, allowing for up to two mismatches. ChIP-seq reads were mapped to the genome as fastq files as described above without the size-filtering step. Read coverage was calculated at each base using bedtools (Quinlan and Hall 2010). Coverage values were normalized by dividing by the number of total mapped reads for each library and enrichment was calculated by dividing the normalized coverage for IP libraries by the normalized coverage of their respective input libraries.

### Telomeric or subtelomeric small RNA analysis on public datasets

Small RNA analysis of public datasets was conducted using the tinyRNA pipeline (Tate et al. 2023). Seven repeats of the telomeric sequence, TTAGGC, and 2 kb of subtelomeric sequences were added as artificial contigs to the *C. elegans* WS279 reference genome and corresponding annotation file to identify telomeric and subtelomeric small RNAs, along with other small RNA classes. Because gene products or transposons are present within the 2 kb subtelomeric regions of chromosome arms IIIR and IVL, these regions were further subdivided into IIIR_short (1–1350 nt), IIIR_long (1351–2000 nt), IVL_short (1–528 nt), and IVL_long (529–2000 nt). Within the tinyRNA pipeline, default parameters were used except that sequences with counts ≤ 1 were removed before alignment with Bowtie, and up to 2 mismatches (end_to_end: 2) were allowed for telomeric small RNAs unless otherwise specified. First, small RNA datasets from N2 embryos, L4s, young adults, and isolated adult gonads were analyzed (Reed et al. 2020; Montgomery et al. 2021; Knittel et al. 2024). To determine potential telomeric and subtelomeric small RNA–associated Argonaute proteins, 21 publicly available Argonaute IP small RNA sequencing datasets were analyzed (Charlesworth et al. 2021; Seroussi et al. 2023), including ALG-1, ALG-2, ALG-3, ALG-4, ALG-5, PRG-1, HRDE-1, NRDE-3, C04F12.1, ERGO-1, WAGO-1, WAGO-4, WAGO-10, PPW-1, PPW-2, RDE-1, SAGO-1, SAGO-2, CSR-1, CSR-1a, and CSR-1b. Small RNA datasets from a subset of Argonaute mutants were analyzed to determine whether small RNAs derived from telomeric repeats and subtelomeric regions are dependent on these Argonautes (Montgomery et al. 2021; Seroussi et al. 2023), including PRG-1, CSR-1, PPW-2, HRDE-1, and WAGO-1. To gain insights into the biogenesis of the telomeric and subtelomeric small RNAs, small RNA datasets from RNAi pathway perturbations (Montgomery et al. 2021; Knittel et al. 2025), including mutants of *rde-1*, *rde-4*, *mut-16*, and *dcr-1* RNAi knockdown, were also analyzed. Further, because perfectly matching small RNAs to the 7-repeat telomeric sequence are rare, analyses were also performed without filtering of sequences with counts ≤ 1 for N2 small RNA samples. Differentially expressed small RNAs between Argonaute IP and input samples, between N2 samples from different developmental stages and N2 adult whole worms, and between mutants and corresponding controls were identified using DESeq2 within the tinyRNA workflow, with a threshold of log2 fold change > 1 and adjusted p-value < 0.05. Results for telomeric and subtelomeric small RNAs are detailed in Supplemental Tables 4-6. R, Adobe Illustrator, and Excel were used for plotting.

### PacBio HiFi sequencing and de novo genome assembly

Genomic DNA was isolated based on the Qiagen genomic DNA purification using Puregene Cell Kit (CAT:158043). DNA concentration and purity were assessed using NanoDrop spectrophotometry and Qubit fluorometric quantification. High-molecular-weight genomic DNA was then submitted to the sequencing facility for PacBio library preparation and SMRT sequencing.

HiFi circular consensus sequencing (CCS) reads generated on the PacBio SMRT platform were obtained as BAM files and converted to FASTQ format prior to downstream analysis. Read quality was evaluated using FastQC to assess sequencing metrics including read length distribution and base quality scores. Processed HiFi reads were used for de novo genome assembly using Hifiasm v0.16 to generate contig assemblies. Genome assembly jobs were executed on the Longleaf high-performance computing cluster at UNC Chapel Hill using SLURM workload manager. The primary contig assembly output generated by Hifiasm was used for all downstream analyses.

Preprocessing and assembly scripts are available on GitHub https://github.com/ambika444/Telomere_Assembly_pipeline.

## Results

### Loss of endogenous sRNAs or XNP-1/ATRX does not promote ALT

Most human cancers achieve immortality by expressing TERT telomerase reverse transcriptase and are termed telomerase-positive cancers (Shay and Wright 2019). High levels of TERRA and loss of the histone H3.3 chaperone subunits ATRX and DAXX commonly occurs in ∼10% of human cancers that maintain their telomeres via the telomerase-independent ALT pathway (Lovejoy et al. 2012; Flynn et al. 2015; Muoio and Fouquerel 2025). The ATRX/DAXX complex mediates histone H3.3 incorporation at telomeres and interacts with methylated H3K9 and HP1 to promote heterochromatin formation (Wong et al. 2010). ATRX has been proposed to repress ALT by protecting telomeres from replication stress and TERRA expression (Udugama et al. 2015). Because human ALT cancers are characterized by ATRX-mediated heterochromatin dysfunction (Heaphy et al. 2011 Jul 22; Lovejoy et al. 2012; Lin et al. 2026), we reasoned that small RNA factors that promote heterochromatin silencing might regulate the establishment of ALT.

We previously studied the Bristol N2 wild type *C. elegans* genome and identified ∼1,200 Interstitial Telomere Sequence (ITS) tracts, which are degenerate telomere sequences that are scattered on chromosome arms and are enriched for heterochromatin marks (Frenk et al. 2019). While telomeric small RNAs with 0-1 mismatches were very rare, abundant telomeric small RNAs with 2-3 mismatches to the *C. elegans* TTAGGC telomere sequence were identified, and some mapped to heterochromatic ITS tracts (Frenk et al. 2019). We previously found that 10-20% of *C. elegans* strains that are deficient for telomerase can be maintained indefinitely through the activation of ALT when maintained by transferring 100-400 larvae per *trt-1* mutant strain once a week (Alt assays) (Cheng et al., 2012). In contrast, ALT never occurs when 4-6 larvae per *trt-1* mutant strain are transferred to fresh plates once per week (Mrt assays) (Cheng et al. 2012). We tested the hypothesis that telomeric small RNAs might repress ALT by passaging *trt-1* telomerase mutant strains that were deficient for biogenesis of either primary or secondary sRNAs using Alt assays. Loss of either primary sRNAs (*trt-1; dcr-1)* or downstream secondary sRNAs (*trt-1; mut-7* and *trt-1; mut-14)* did not significantly alter the proportion of strains that survive via ALT (Figure 1A, Table S1). Similarly, loss of the *C. elegans* ortholog of the histone H3.3 chaperone ATRX, XNP-1, itself did not affect the establishment or maintenance of ALT (Figure 1A, Table S1). Although most small RNA or XNP-1 deficient mutants displayed a faster onset of sterility, the *xnp-1* deletion *ok1823* repressed sterility during maintenance in Alt assays in comparison to the *xnp-1 tm678* deletion. This effect on the onset of sterility in Alt assays could be caused by a mutation linked to these deletions, which were created using a mutagen that peppers the genome with mutations (Moerman and Barstead 2008). A future study could ask if the *xnp-1(ok1823)* deletion or a mutation linked to this deletion is responsible for the slower onset of sterility of *trt-1; xnp-1(ok1823)* mutants in Alt assays. Overall, our Alt assays analysis reveals that neither loss of sRNA-mediated genome silencing nor XNP-1/ATRX-mediated heterochromatin formation is sufficient to promote ALT in *C. elegans* nematodes.

**Figure 1:**
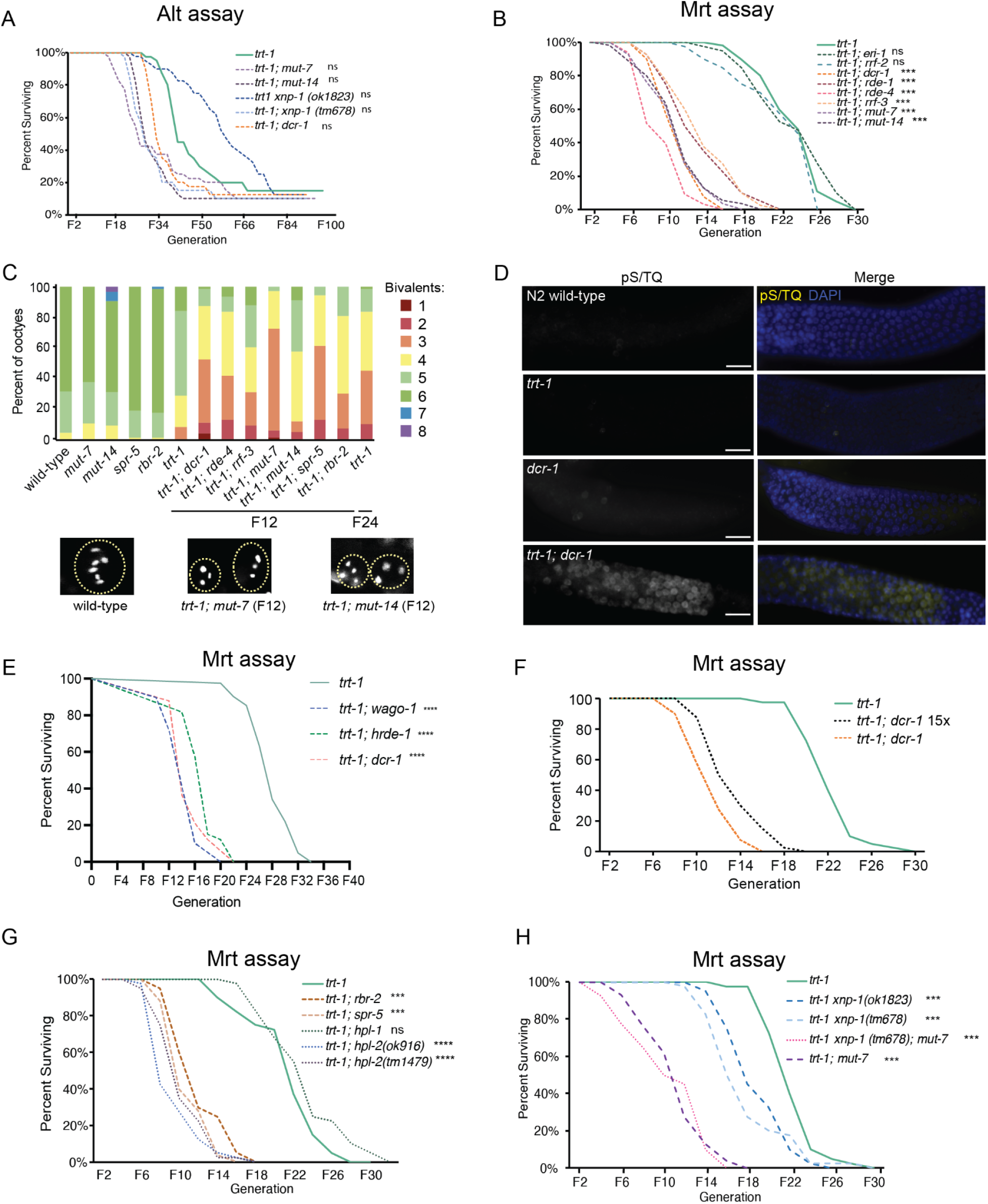
Endogenous small RNA and chromatin factors repress telomere stability in the absence of telomerase. (A) Survival curve of telomerase-deficient strains passaged at ∼100 worms weekly for 40 independent lines. One allele of *xnp-1(ok1823)* extended survival but did not increase survival in Alt assays (Table S1). The number of survivors was compared between the *trt-1* and double mutant groups. Statistical analysis was performed, and the detailed results are provided in Supplementary Table 1. (B) Survival curve of primary or secondary sRNA deficient telomerase mutants when 6 larval L1s are passaged weekly. (C) Small RNA and heterochromatin mutant *trt-1* lines have high levels of chromosome fusions, comparable to *trt-1* single mutants at F24. Fusions were assayed by diamidino-2-phenylindole (DAPI) staining of oocytes in diakinesis. Each bivalent in wildtype represents a single paired chromosome. Representative images of DAPIstained oocyte nuclei are shown in panels below the graph. (D) High levels of phospho(Ser/Thr) staining in the meiotic gonad of *trt-1; dcr-1* at F6 compared to N2 or *trt-1*. Scale bar, 25 um. (E) Loss of HRDE-1 or WAGO-1 Argonaute proteins mimics the accelerated sterility observed in a *trt-1; dcr-1* mutant background in Mrt assays. (F) A *trt-1; dcr-1* strain was crossed with *dcr-1* males fifteen times to yield a *trt-1; dcr-1* 15x strain that displayed the same accelerated sterility observed for the initial *trt-1; dcr-1* double mutants in Mrt assays. (G) Survival curve of double mutants for *trt-1* and chromatin factors passaged by 6 L1s weekly. (H) Mrt assays reveal that histone H3.3 remodeling protein XNP-1 represses senescence in *trt-1* mutants epistatically to the small RNA factor MUT-7. (B), (F)(G)(H) ***p < 0.001; ****p < 0.0001; ns, not significant, (Log rank test, n=40 independent lines for each genotype). (E) *trt-1; wago-1* = 39 lines per genotype. *trt-1; hrde-1* and *trt-1; dcr-1* = 33 lines per genotype.

### Small RNA factors repress rapid sterility when telomerase is deficient

*C. elegans trt-1* telomerase mutant strains uniformly develop sterility in response to telomere erosion after 20-30 generations of growth when 6 L1 larvae are passaged weekly in Mrt assays (Ahmed and Hodgkin 2000; Meier et al. 2006; Cheng et al. 2012). The sterility of late-generation *trt-1* mutants is induced by telomere fusions (Cleal and Baird 2020). Intriguingly, we found that *trt-1* mutants that lack some primary or secondary sRNA biogenesis factors become sterile significantly faster than *trt-1* single mutants in Mrt assays (Figure 1B, Table S2). Wild type *C. elegans* strains contain a haploid complement of 6 chromosomes that pair during meiosis and chromosome number can be detected in mature oocytes that are arrested in meiosis I as 6 DAPI-positive spots. Telomere fusions are detected when fewer than 6 DAPI-positive spots are present in oocytes of late-generation telomerase mutant strains. *trt-1* mutant strains that lack some genes that promote small RNA biogenesis or function rapidly accumulated end-to-end chromosome fusions by generation F12, a characteristic of telomerase mutants that have grown for more than 20 generations (Figure 1C).

*C. elegans* chromosome fusions cause non-disjunction events during meiosis that elicit dominant embryonic lethality or X0 male phenotypes in progeny of heterozygous F1 animals. We asked if early generation *trt-1; dcr-1* double mutant animals possessed chromosome fusions by performing genetic crosses that would be capable of revealing a single heterozygous end-to-end chromosome fusion (Ahmed and Hodgkin, 2000). We crossed four independently derived *trt-1 dcr-1* F4 hermaphrodites with N2 wild-type males but failed to detect fused chromosomes in F1 progeny of F4 mothers in contrast to F1 progeny of F24 *trt-1* mutant controls (p = 3.841e-05 for males and p = 3.61e-05 for dead embryos, Mann Whitney U test) (Table S3). However, early-generation *trt-1; dcr-1* mutant strains displayed high levels of DNA damage signaling that were not observed for *trt-1* and *dcr-1* single mutant controls (Figure 1D). Together, these results indicate that loss of small RNAs and telomerase elicits telomere damage but not telomere fusions in early generations. The rapid sterility of these strains in later generations is promoted by abundant end-to-end chromosome fusions.

We created a number of sRNA-deficient *trt-1* double mutants to define additional small RNA factors that elicit rapid sterility in the absence of telomerase. Rapid sterility in the absence of telomerase occurred upon loss of the Dicer nuclease DCR-1 and by RDE-4, which binds endogenous dsRNA and forms a complex with DCR-1, and the Argonaute RDE-1, which acts with the RNA-dependent RNA polymerase RRF-3 to promote production of dsRNA-derived primary sRNAs (Tabara et al. 2002; Steiner et al. 2009; Gent et al. 2010) (Figure 1B). Loss of the MUTATOR proteins MUT-7 and MUT-14, which act in germ granules to create 22G secondary sRNAs that can promote transcriptional silencing in the nucleus (Ketting et al. 1999; Tijsterman et al. 2002; Billi et al. 2018), caused rapid sterility of *trt-1* mutants (Figure 1B). Two other small RNA factors, RRF-2 and ERI-1, did not affect *trt-1* mutant sterility (Figure 1B), indicating that the endogenous 26G sRNA pathway that requires ERI-1 (Pavelec et al. 2009; Gent et al. 2010) is dispensable for telomere stability in the absence of telomerase.

We crossed *dcr-1* and *mut-7* single mutants with wild-type males, singled F2 progeny that were homozygous mutants for *dcr-1* or *mut-7*, and isolated genomic DNA from F4, F6 and F8 generation descendants. Southern blotting revealed discrete telomere bands that display modest telomere erosion for freshly homozygous *dcr-1* and *mut-7* mutant telomeres (Figure S1A, B), although none of the small RNA single mutants we tested became progressively sterile or accumulated chromosome fusions (Figure 1C). This indicates that although telomere erosion can occur in RNAi-deficient *C. elegans* mutants, that bouts of telomerase activity will occasionally lengthen telomeres, although we did not test this experimentally.

We asked if the accelerated sterility of small RNA-deficient *trt-1* mutant strains might be explained by short telomeres inherited from gametes derived from small RNA-deficient strains by crossing a *trt-1; dcr-1* double mutant hermaphrodite with *dcr-1* mutant males, singling *trt-1 +/−; dcr-1 −/−* F1 hermaphrodites and isolating *trt-1; dcr-1* double mutant F2. This cross was repeated 15 times to create a *trt-1; dcr-1* double mutant strain whose telomeres were exclusively derived from the *dcr-1* mutant background. The *trt-1; dcr-1* 15x strain became sterile slightly slower than freshly created *trt-1; dcr-1* double mutants, although this difference was not statistically significant (p = 0.07) (Figure 1E and 1F). These results indicate that telomeres transmitted from the *dcr-1* mutant are of normal length and cannot explain the rapid sterility of *trt-1; dcr-1* double mutants. Therefore, telomerase likely heals telomeric damage that occurs in *dcr-1(mg375)* mutants.

We analyzed the dynamics of telomere erosion in sRNA-deficient *trt-1* mutant strains using terminal restriction fragment analysis but did not observe a significant difference in the rate of telomere erosion when passaging multiple worms per plate, in comparison to *trt-1* single mutant controls (p > 0.7 in both cases) (Figure S1A, B). However, when we singled individual worms for each genotype, we observed significant levels of telomere instability in small RNA-deficient *trt-1* mutant strains in comparison to *trt-1* single mutant controls. *trt-1; mut-7* and *trt-1; dcr-1* mutants had a significantly larger variance in the rate of telomere erosion than *trt-1* mutants, which resulted from atypical shortening and elongation events (Figure S2A, B). We hypothesize that the atypical telomere shortening events cause accelerated sterility.

### Heterochromatin factors repress rapid sterility when telomerase is deficient

We considered that small RNAs might repress rapid sterility by directing chromatin factors to telomeres to promote deposition of heterochromatic marks. Chromatin immunoprecipitation experiments have previously revealed that *C. elegans* telomeres are enriched for the H3K9me2 heterochromatin mark. However, telomeres were not enriched for H3K9me3, which is bound by another *C. elegans* HP1 homolog HPL-1 (Zeller et al. 2016; McMurchy et al. 2017). Similar to primary and secondary small RNA-deficient *trt-1* strains, loss of HPL-2, which interacts with H3K9me2, but not HPL-1 caused accelerated senescence in *trt-1* mutants (Figure 1G). H3K4 methylation, which is associated with regions of active transcription, must be removed by demethylases prior to deposition of repressive H3K9 methylation (Greer et al. 2011; Li and Kelly 2011). We found that loss of either of the two *C. elegans* H3K4 demethylases RBR-2/KDM5 or SPR-5/LSD1 caused rapid sterility of *trt-1* mutants (Figure 1G). Thus, proteins that promote H3K4 demethylation and H3K9me2 deposition prevent telomere instability in the absence of telomerase.

The histone remodeling protein XNP-1/ATRX has previously been implicated in telomeric heterochromatin formation and interacts with unmethylated H3K4 and methylated H3K9 deposited on the histone variant H3.3 (Villard et al. 1999; Bender et al. 2004; Cardoso et al. 2005). *trt-1 xnp-1* double mutants became sterile significantly faster than *trt-1* single mutants but not as fast as small RNA deficient *trt-1; mut-7* double mutants (Figure 1H). Additionally, as loss of both *xnp-1* and *mut-7* did not exacerbate the accelerated sterility phenotype of sRNA-deficient *trt-1* mutant strains (Figure 1H), these genes likely function in the same telomere stability pathway. These data suggest that the chromatin remodeler XNP-1/ATRX acts in germ cells to promote heterochromatin formation at telomeres in response to small RNAs. As XNP-1 modifies H3.3, small RNA-dependent methylation of distinct histones, like canonical histone H3, may contribute to telomere stability in parallel with H3.3.

### Increased expression of TERRA in small RNA and chromatin deficient *trt-1* strains

Small RNAs and chromatin factors have been reported to promote silencing of TERRA (Vrbsky et al. 2010; Rossiello et al. 2017), so we measured the abundance of TERRA by qRT-PCR at three chromosome ends using primers within their H3K9me2-coated subtelomere domains (Figure 2A-C). We confirmed that H3K9me2 but not H3K9me3 is enriched at *C. elegans* telomeres using high throughput sequencing data sets generated from chromatin immunoprecipitation (ChIP) experiments performed by independent groups (Zeller et al. 2016; McMurchy et al. 2017). Moreover, we observed small H3K9me2 domains that extended into the subtelomere at all *C. elegans* chromosome ends (Figure 2A-C). Compared to *trt-1* single mutant controls, TERRA was upregulated in early-generations in *trt-1; rde-4*, *trt-1; mut-14* and *trt-1; spr-5* double mutants, but not in *trt-1* or small RNA-deficient single mutant controls (Figure 3A). These results are distinct from previous observations that TERRA transcription can be specifically induced at damaged or shortened telomeres of mammalian cells that possess telomerase (Porro et al. 2014; Rossiello et al. 2017).

**Figure 2:**
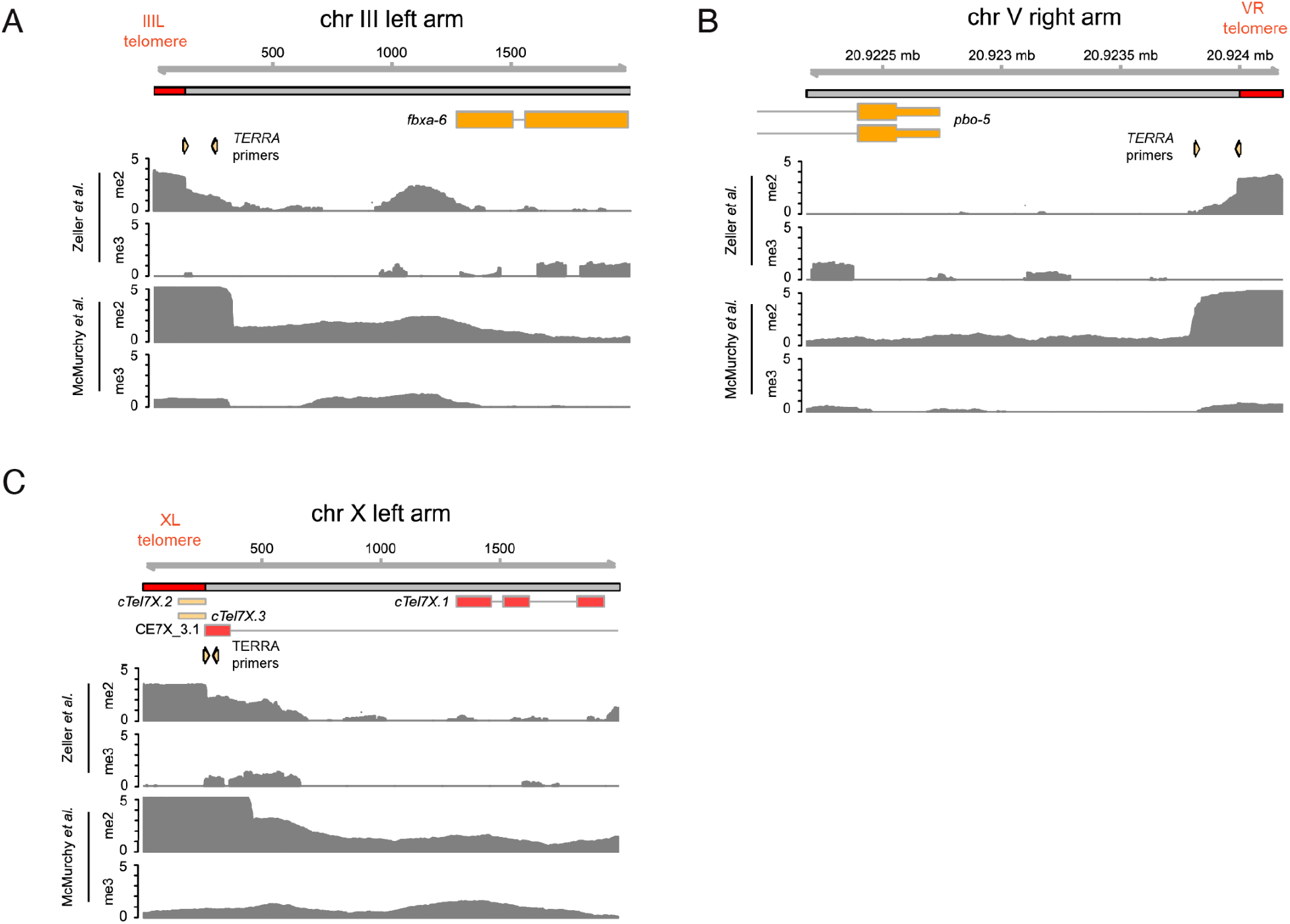
Analysis of telomeric chromatin and small RNAs in published datasets. (A)-(C) ChIP-seq enrichment profiles at three chromosome ends constructed using published data (McMurchy et al., 2017, Zeller et al., 2016). Small orange arrows between gene models and data tracks show the position of primers used in TERRA qRT-PCR experiments.

**Figure 3:**
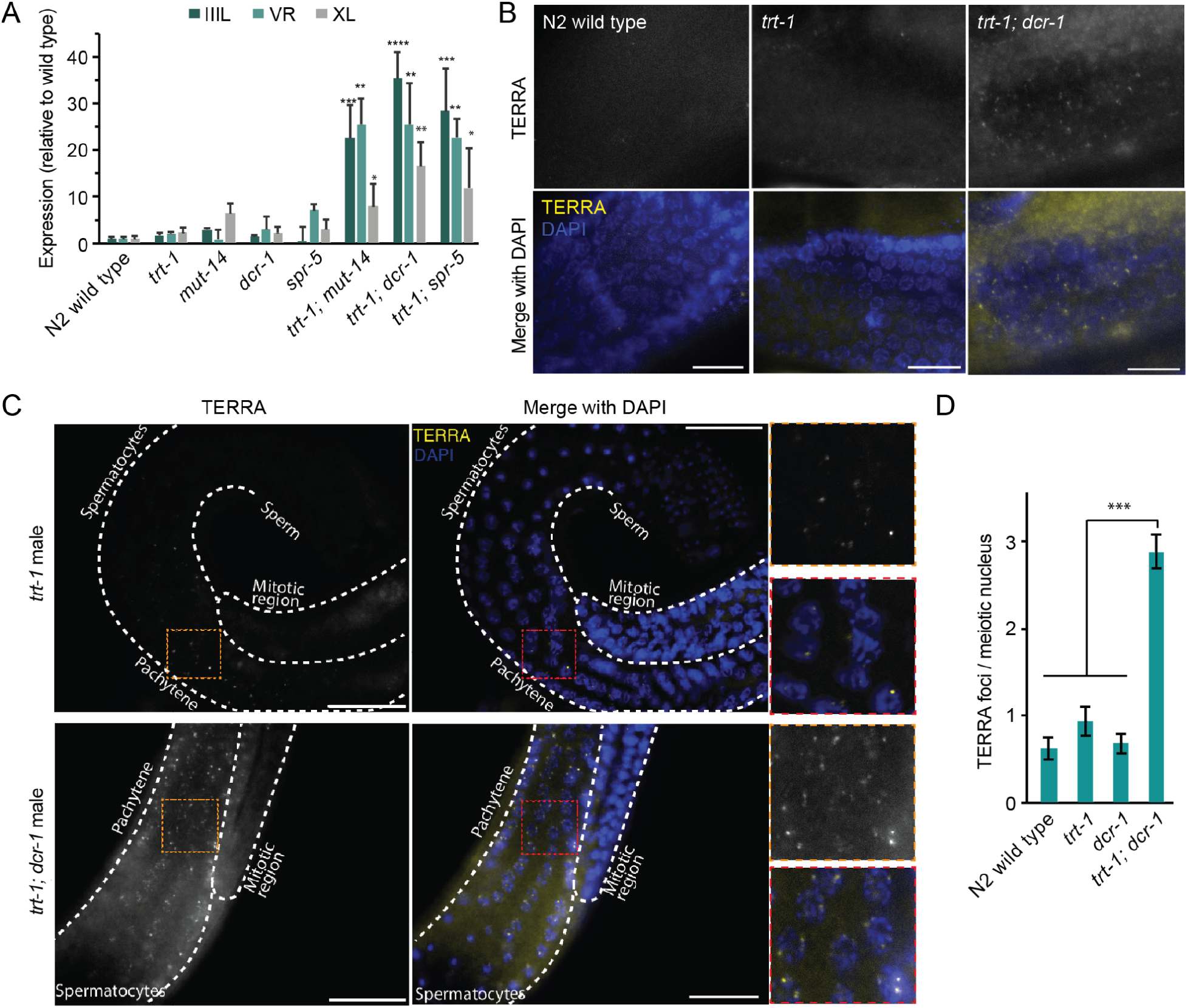
(A) Expression of TERRA from subtelomeres *IIIL*, *VR*, and *XL* is increased in sRNA and heterochromatin deficient telomerase mutants. cDNA was reverse transcribed with a telomere specific primer and qPCR was performed using subtelomere-specific primer pairs and normalized to *act-1*. Expression data are plotted relative to wildtype and represent the mean ± SEM of three biological replicates. Comparison between expression level of wild type and mutants were made using unpaired-T test. (B) RNA FISH signals of TERRA in meiotic germlines of hermaphrodites of the indicated genotypes. TERRA foci are present in wildtype and *trt-1* in the pachytene region and increased in *trt-1; dcr-1.* Scale bar, 10um. (C) Representative RNA FISH images of TERRA in males show foci in the pachytene region of the germline which are increased in *trt-1; dcr-1* compared to *trt-1.* Scale bar, 20 um. (D) Quantification of germline TERRA foci (mean ± SEM, n=10 germlines).

We next used RNA Fluorescence *In Situ* Hybridization (FISH) to confirm the effects on TERRA expression and to examine its cell and tissue localization. In wild-type control and *trt-1* mutant hermaphrodite and male worms, TERRA was expressed at low levels in the soma as well as in the pachytene region of the meiotic germline (Figure 3B-D, S3-4), which corresponds to the location of several lncRNA-mediated germline silencing processes (Taylor et al. 2015), including Meiotic Silencing of Sex Chromosomes in mammals and *C. elegans (*Turner 2015). Compared to both wild-type control and *trt-1* or *dcr-1* single mutant strains, *trt-1; dcr-1* double mutant strains had increased expression of TERRA in the germline (Figure 3B-D, S4) and soma (Figure S3-4) by the F6 generation. We observed an increase in the number of TERRA foci per nucleus in sRNA-deficient *trt-1* mutants, which might be telomeres (Figure 3D). We also observed TERRA expression by RNA FISH in the cytoplasm and nucleoplasm of sRNA-deficient *trt-1* cells (Figure 3B-C, S4), consistent with possible non-telomeric functions of TERRA, which could be epigenetic in nature (Chu et al. 2017; Roake and Artandi 2020) or with a role for TERRA in protecting telomeres *in trans* (Montero et al. 2018). Therefore, when telomerase is absent, endogenous small RNAs either repress telomere transcription or promote TERRA degradation, possibly in the context of DNA damage signaling events (Figure 1D).

### Subtelomeric small RNAs associate with a subset of Argonaute proteins

Telomeric sRNAs have been observed in protists, plants, and mammals (Cao et al. 2009; Couvillion et al. 2009; Vrbsky et al. 2010). We previously interrogated 92 publicly available small RNA sequencing libraries created from RNA isolated from various *C. elegans* strains and found rare reads that mapped perfectly to the *C. elegans* telomere repeat (TTAGGC)_n_, occurring at a frequency of ∼0.1 read per million (Frenk et al. 2019). piRNAs with up to 3 mismatches can target regions of the *C. elegans* genome for silencing (Bagijn et al. 2012), and we found that telomeric sRNAs with up to 3 mismatches were substantially more abundant than those with perfect telomere repeat sequence (∼10 reads per million) (Frenk et al. 2019). In this study, we examined small RNAs from wild type at different developmental stages for telomeric and subtelomeric small RNAs. Perfect telomeric small RNAs are present in wild type adults or embryos, but not in dissected germlines or L4 larvae (Figure S5A). As wild type adults contain embryos, this implies that rare perfect telomeric small RNAs are created in embryos. On the other hand, telomeric small RNAs with up to 2 mismatches were present in all samples, including dissected adult germlines (Table 1 and Figure S5A). Dissected wild type germlines were enriched with small RNAs targeting IR in comparison to wild type adults. Next, we compared the subtelomeric small RNAs profiles of wild type L4 stage and adult samples. Small RNAs targeting IR, IIIL, IIIR, IVL, IVR, and VL are more enriched in L4 samples, while IIL and IIIR are enriched in adult samples. Subtelomeric small RNA levels were not different when embryos were compared with wild type adults.

**Table 1.** Summary of differentially expressed telomeric or subtelomeric small RNAs in N2 samples and Argonaute IP small RNA datasets.

| Features | Wild type |  |  | Argonaute IP vs input |  |  |  |  |  |  |  |  |  |  |  |
|---|---|---|---|---|---|---|---|---|---|---|---|---|---|---|---|
|  | N2 L4 whole vs Adult whole | N2 Adult gonad vs Adult whole worm | N2 Embryos vs adult whole worm | HRDE-1 | WAGO-1 | PPW-1 | PPW-2 | RDE-1 | CSR-1 | CSR-1a | CSR-1b | PRG-1 | WAGO-4 | ALG-1 | ALG-2 |
| telomere_mismatchs antisense |  |  |  | ++ |  |  | ++ |  | -- |  |  |  |  |  |  |
| telomere_mismatchs sense |  |  |  | ++ | ++ | ++ |  |  |  |  |  |  |  |  |  |
| IL subtelo antisense |  |  |  | + | + |  | ++ | -- | -- |  | -- | ++ |  |  | -- |
| IL subtelo sense |  |  |  | + | + |  | ++ |  | -- |  | -- | -- | - |  | -- |
| IR subtelo antisense |  | ++ |  |  | + |  |  | + | + | ++ | ++ |  |  | -- | -- |
| IR subtelo sense | + | ++ |  | -- | -- | -- | -- | + | -- |  |  | -- | -- |  |  |
| IIL subtelo antisense |  |  |  | ++ |  | ++ | ++ |  | -- |  |  |  |  |  | -- |
| IIL subtelo sense | - |  |  | ++ | + |  | -- |  | -- |  |  | -- | -- |  |  |
| IIR subtelo antisense |  |  |  |  |  |  |  |  |  |  |  |  |  |  |  |
| IIR subtelo sense |  |  |  |  |  |  |  |  |  |  |  |  |  |  |  |
| IIIL subtelo antisense | + |  |  |  |  |  | ++ |  | - |  |  |  |  |  |  |
| IIIL subtelo sense |  |  |  |  |  |  |  |  |  |  |  |  |  |  |  |
| IIIR long subtelo antisense | - |  |  |  |  |  |  |  |  |  |  |  |  |  |  |
| IIIR long subtelo sense |  |  |  |  |  |  | ++ |  |  |  |  |  |  |  |  |
| IIIR short subtelo antisense | - |  |  |  | ++ |  |  |  |  |  |  |  | ++ |  |  |
| IIIR short subtelo sense | + |  |  |  |  |  | ++ |  |  |  |  |  |  |  |  |
| IVL long subtelo antisense |  |  |  |  |  |  |  |  |  |  |  |  |  |  |  |
| IVL long subtelo sense | ++ |  |  |  |  |  |  |  |  |  |  |  |  |  |  |
| IVL short subtelo antisense |  |  |  |  |  |  |  |  |  |  |  |  |  |  |  |
| IVL short subtelo sense |  |  |  |  |  |  |  |  |  |  |  |  |  |  |  |
| IVR subtelo antisense | ++ |  |  |  |  |  |  |  |  |  |  |  |  |  |  |
| IVR subtelo sense | ++ |  |  | ++ |  |  |  |  |  |  |  |  |  |  |  |
| VLsubtelo antisense |  |  |  |  |  |  | ++ |  | -- |  |  |  |  |  |  |
| VL subtelo sense | ++ |  |  |  |  |  | ++ |  | -- |  |  |  |  |  |  |
| VR subtelo antisense |  |  |  |  |  |  |  |  |  |  |  |  |  |  |  |
| VR subtelo sense |  |  |  |  |  |  | ++ |  |  |  |  |  |  |  |  |
| XL subtelo antisense |  |  |  |  |  |  |  |  |  |  |  |  |  |  |  |
| XL subtelo sense |  |  |  | + |  |  | ++ |  | -- |  | -- |  |  | -- | -- |
| XR subtelo antisense |  |  |  |  |  |  |  |  |  |  |  |  |  |  |  |
| XR subtelo sense |  |  |  |  |  |  |  |  |  |  |  |  |  |  |  |

Symbols indicate differential expression analysis using DESeq2 from tinyRNA. "+"/"-" denote log2 fold change > 1 with p-adjusted < 0.05, and "++"/"--" denote log2 fold change > 2 with p-adjusted < 0.05. Up to 2 mismatches were allowed for mapping to telomeric repeats during the analysis. Details are included in Table S4-6.

We compared levels of telomeric small RNAs with mismatches from wild type with those of mutants for small RNA genes that promote telomere stability in the absence of telomerase, using data sets for *dcr-1*, *rde-4*, *mut-7*, and *mut-14* mutants, and found that levels of telomeric sRNAs were sometimes depleted in these backgrounds, but not always (Table 2). We also interrogated older data sets of small RNAs that immunoprecipitate with Argonaute proteins that are known to promote silencing in the germline, HRDE-1 and WAGO-1, as well as with the germline anti-silencing Argonaute CSR-1 (Gu et al., 2009, Claycomb et al., 2009, Ashe et al., 2012). Enrichment of small RNAs for each protein was calculated by dividing the number of reads in the immunoprecipitate library by the number of reads in the corresponding input library. Telomeric small RNAs were enriched in WAGO-1 immunoprecipitates but not for HRDE-1 or CSR-1 (Figure S5B). Enrichment of subtelomeric small RNAs in WAGO-1 and HRDE-1 was detected for most chromosome ends (Figure S5C). Interestingly, CSR-1 was only enriched for subtelomeric small RNAs corresponding to rDNA at the right end of Chromosome I (Wicky et al., 1996). It is therefore possible that CSR-1 takes a predominantly anti-silencing function at the *IR* subtelomere.

**Table 2.** Summary of differentially expressed telomeric or subtelomeric small RNAs with disruption of a subset of small RNA pathway genes.

| Features | Mutant vs WT |  |  |  |  |  |  |  |  |  |
|---|---|---|---|---|---|---|---|---|---|---|
|  | mut-16(pk710) gonads vs N2 gonads | hrde-1 (tm1200) vs N2 | wago-1(tm1414) vs N2 | ppw-2 (tm1120) vs N2 | rde-1 (ne219) vs N2 | rde-4(sy6665) L4 vs N2 L4 | dcr-1RNAi vs L4440 | csr-1 (tm892) vs N2 | prg-1(tm872) vs N2 | prg-1(n4357) gonads vs N2 gonads |
| telomere_mismatchs antisense | -- | -- |  | -- |  |  |  | -- |  | + |
| telomere_mismatchs sense |  |  |  | -- |  |  |  |  | ++ | + |
| IL subtelo antisense | -- | - |  |  |  |  |  | + | -- | -- |
| IL subtelo sense | -- | - | - |  |  |  |  | + | -- | -- |
| IR subtelo antisense |  |  |  | - |  |  |  | ++ | + | ++ |
| IR subtelo sense |  |  |  |  |  |  |  |  |  |  |
| IIL subtelo antisense | -- | -- |  | -- | + | + |  |  | ++ | ++ |
| IIL subtelo sense | -- | -- |  |  |  |  |  | -- | -- | + |
| IIR subtelo antisense |  |  |  |  |  |  |  |  |  |  |
| IIR subtelo sense |  |  |  |  |  |  |  |  |  |  |
| IIIL subtelo antisense | -- | -- | -- | -- |  |  |  | -- |  | + |
| IIIL subtelo sense |  |  |  |  |  |  |  |  |  |  |
| IIIR long subtelo antisense | -- |  |  |  |  | - |  |  | -- | -- |
| IIIR long subtelo sense | -- |  |  |  |  |  |  |  | -- |  |
| IIIR short subtelo antisense |  |  |  |  |  |  |  |  |  |  |
| IIIR short subtelo sense |  |  | -- |  |  |  |  |  | - | -- |
| IVL long subtelo antisense |  |  |  |  |  |  |  |  | -- |  |
| IVL long subtelo sense |  |  |  |  |  |  |  |  |  |  |
| IVL short subtelo antisense |  |  |  |  |  |  |  |  |  |  |
| IVL short subtelo sense |  |  |  |  |  |  |  |  |  |  |
| IVR subtelo antisense | -- |  |  |  | ++ | + |  |  | ++ | ++ |
| IVR subtelo sense | -- |  |  |  | ++ | + |  |  | ++ | ++ |
| VLsubtelo antisense |  |  |  | - |  |  |  | ++ | - |  |
| VL subtelo sense |  |  |  | - |  |  |  | + | - |  |
| VR subtelo antisense |  |  |  |  |  |  |  |  |  |  |
| VR subtelo sense |  |  |  |  |  |  |  |  |  |  |
| XL subtelo antisense |  |  |  |  |  |  |  |  |  |  |
| XL subtelo sense | -- | + | -- |  |  |  |  |  | -- | -- |
| XR subtelo antisense |  |  |  |  |  |  |  |  |  |  |
| XR subtelo sense |  |  |  |  |  |  |  |  |  |  |

Symbols indicate differential expression analysis using DESeq2 from tinyRNA. "+"/"-" denote log2 fold change > 1 with p-adjusted < 0.05, and "++"/"--" denote log2 fold change > 2 with p-adjusted < 0.05. Up to 2 mismatches were allowed for mapping to telomeric repeats during the analysis. Details are included in Table S4-6.

We next interrogated comprehensive Argonaute immunoprecipitation small RNA datasets with biological replicates to identify the specific Argonaute proteins that interact with telomeric and subtelomeric small RNAs (Seroussi et al. 2023). As perfect telomeric sRNAs are exceptionally rare, we asked if telomeric sRNAs with up to 2 mismatches associated with Argonaute proteins. We also studied these data sets for subtelomeric sRNAs that uniquely match sequences within 2 kb of each telomere. We used shorter subtelomere segments for *IIIR* and *IVL*, as a *Tc6* transposon that is targeted by sRNAs (*IIIR*) or part of the *frm-5.2* gene (*IVL*) are located within 2 kb of these telomeres. We observed enrichment of small RNAs for multiple subtelomeres over input for three Argonaute proteins: HRDE-1 (n=4), WAGO-1 (n=4), PPW-2 (n=7), C04F12.1 (n=2), NRDE-3 (n=2), SAGO-2 (n=3). In addition, telomeric sRNAs with up to 2 mismatches were enriched for HRDE-1 (sense and antisence), WAGO-1 (sense), PPW-1 (sense) and PPW-2 (antisense) and NRDE-3 (antisense) (Table 1 and Figure S6).

Because HRDE-1, WAGO-1, and PPW-2 showed the strongest enrichment for telomeric or subtelomeric small RNAs, we asked whether these small RNAs depend on these Argonaute proteins by analyzing mutant datasets (Seroussi et al. 2023). We studied telomeric and subtelomeric sRNAs and observed that *hrde-1* mutants were significantly depleted for subtelomeric IL and IIL sRNAs (sense and antisense), and IIIL sRNAs (antisense) (Table 2). *wago-1* mutants were significantly depleted for 3 sense (IL, IIIR_short, and XL) and 1 antisense (IIIL) subtelomere sRNA classes. *wago-1; trt-1* and *trt-1; hrde-1* double mutants both displayed rapid sterility phenotypes in comparison to *trt-1* single mutants (p < 1e-11 and p < 1e-12, respectively, log rank test) (Figure 1E). The depletion of telomeric or subtelomeric small RNAs in *wago-1* and *hrde-1* mutants further support the role of WAGO-1 and HRDE-1 to promote telomere maintenance in the absence of telomerase.

As PPW-2 displayed strong association with telomeric and subtelomeric small RNAs, and four subtelomeres and telomeres were depleted in *ppw-2* mutants. MUT-7 and MUT-14 promote telomere stability in the absence of telomerase (Figure 1B), they function with MUT-16 to promote small RNA amplification in germ granules (Zhang et al. 2011; Uebel et al. 2018). We compared small RNAs from dissected germlines of *mut-16* mutants with wild type and observed significant depletion for small RNAs to IL, IIL, IIIR_long and IVR subtelomeres (sense and antisense), for sRNAs to IIIL (antisense), and to XL (sense) (Table 2).

As loss of the dsRNA-binding protein RDE-4 caused telomere instability in the absence of telomerase (Figure 1B), we studied small RNAs from *rde-4(syb6665)* mutants but found that neither unique sub-telomeric nor telomeric sRNAs were altered (Table 2) (Knittel et al. 2025). RDE-4 interacts with DCR-1, which is an essential nuclease that promotes microRNA biogenesis (Ketting et al. 2001; Knight and Bass 2001; Tabara et al. 2002). We also found that worms depleted for DCR-1 by RNAi knockdown did not display altered telomeric or subtelomeric sRNAs. Therefore, DCR-1 and RDE-4 promote telomere stability in the absence of telomerase indirectly.

Although the CSR-1 anti-silencing Argonaute only interacted with ribosomal small RNA at the IR subtelomere (Table 1), we studied small RNAs from *csr-1* deletion mutants and observed increased levels of subtelomeric small RNAs at 3 subtelomeres, including IL, IR, and VL, loss of telomeric small RNAs at 2 subtelomeres, as well as depleted telomeric small RNAs with mismatches (Table 2). Similarly, the PRG-1 Piwi Argonaute protein only interacted with small RNAs from the IL subtelomere, but *prg-1* deletion mutants displayed 2– to 16-fold increased levels of telomeric small RNAs and subtelomeric small RNAs for 3 subtelomeres (IR, IIL and IVR), as well as 3– to 7-fold decreased levels of subtelomeric small RNAs for 6 subtelomeres (IL, IIL, IIIR, IVL, VL and XL) (Table 2). Therefore, the PRG-1 Piwi protein that interacts with thousands of piRNAs to scan the genome for intruders (Ashe et al. 2012; Bagijn et al. 2012), and the CSR-1 anti-silencing Argonaute protein (Wedeles et al. 2014) affect both telomeric and subtelomeric small RNA levels, but most of these effects are indirect. We did not assess how loss of *csr-1* or *prg-1* affects telomere stability in the absence of telomerase.

We focused on HRDE-1, WAGO-1, and PPW-2 IP small RNA datasets to examine the length and 5’ nucleotide composition of telomeric, subtelomeric, and all small RNA species (Figure 4A-C). Telomeric small RNAs displayed a dominant enrichment at 22 nt and exhibited a strong 5’ G bias (Figure 4A), as shown for all small RNA species as well (Figure 4C), consistent with the characteristic features of endogenous 22G-sRNAs. In contrast, in addition to the 22G-RNA enrichment, subtelomeric small RNAs are also enriched at 31–35 nt in length. Particularly, 33 nt sRNAs with 5’ C bias account for the largest portion of sequencing reads for HRDE-1 and PPW-2. The long sRNAs (31-35nt) all correspond to the sense strand of the 3’ end of the 26s/28s rRNA, *rrn-3.1* (Figure 4D and 5E), which is close to the IR subtelomere. The 3’ end of *rrn-3.1* overlaps with the 5’ end of *F31C3.14* (Figure 4E), which may contribute to creation of an unusually long class of sRNAs that interact with Argonaute proteins that normally strongly prefer 22 nucleotide RNAs (Figure 4C). No genes are present within 2 kb of other subtelomeres, except for IVL, which is why the ‘IVL short’ subtelomere that lacks the 5’ end of the gene *frm-5.2* was chosen for small RNA analysis.

**Figure 4:**
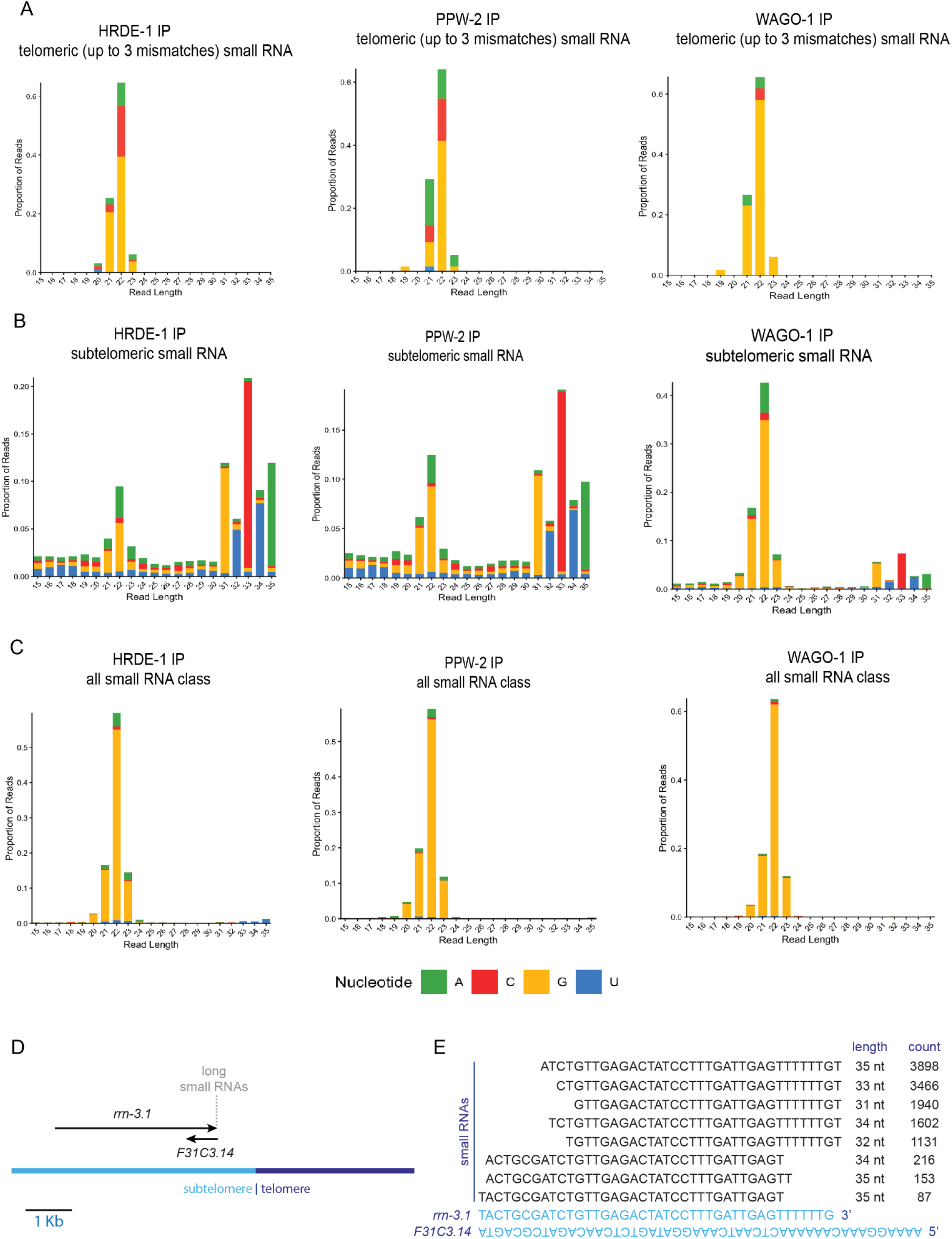
Length and 5’ nucleotide distribution of telomeric and subtelomeric small RNA reads from PPW-2, HRDE-1, WAGO-1 IP small RNA sequencing datasets. (A) Distribution of telomeric small RNAs. Up to 3 mismatches were allowed to map reads to seven telomere repeats (TTAGGC). (B) Distribution of subtelomeric small RNAs. (C) Distribution of small RNAs across all classes. The average distribution from two biological replicates is shown. (D) Diagram showing the position of ribosomal 26s/28s rRNA(*rrn-3.1*) at IR telomere and subtelomere. (E) Abundant long small RNAs overlapping with the end of *rrn-3.1*. 5’ and 3’ ends of *F31C3.14* and *rrn-3.1*, respectively, are shown.

**Figure 5:**
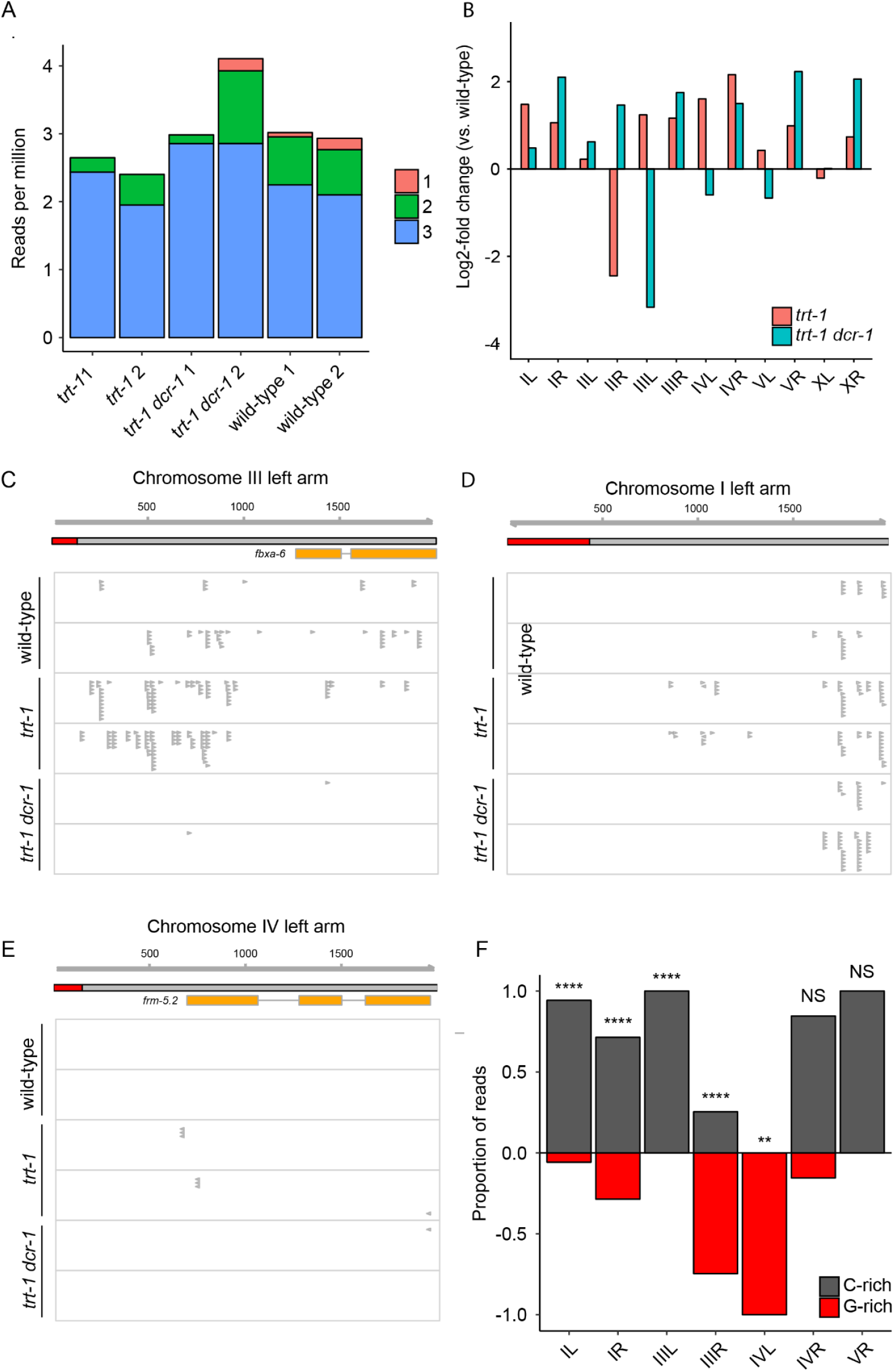
Upregulation of subtelomeric sRNAs in telomerase mutants. (A) Number of telomere-mapping reads in wild-type, *trt-1,* and *trt-1;dcr-1* libraries. Numbers 1-3 indicate the number of mismatches. (B) Log2fold change in sRNA abundance at subtelomeres relative to wildtype in *trt-1* and *trt-1; dcr-1* animals. (C)-(E) 22G sRNAs at three subtelomeres. (F) Proportion of subtelomeric sRNAs mapping to the strand corresponding to the G-rich (TTAGGC) and C-rich (TAAGCC) telomere strand in *trt-1* animals. **** p < 0.0001; ** p < 0.01; ns, not significant (Binomial test with Bonferroni correction). The p values refer to the probability of observing the given proportion of G-rich or C-rich telomeric reads at each chromosome end under the null hypothesis that G– and C-rich reads are present in equal proportions.

### Small RNAs at subtelomeres altered in the absence of telomerase

We asked if loss of telomerase might alter levels of telomeric small RNAs in a manner that might depend on Dicer by isolating RNA from wild-type control, *trt-1* single mutant, and *trt-1; dcr-1* double mutant strains and preparing small RNA libraries. We sequenced these libraries deeply in an effort to detect rare perfect telomeric sRNAs. We obtained ∼20,000,000 reads each for 6 data sets and found that our *trt-1; dcr-1* data sets were strongly depleted of 22G RNAs that were previously reported to depend on *dcr-1* (Figure S7A) (Gent et al. 2010). Loss of TERT promotes stem cell proliferation in mice, due to a non-telomeric function of telomerase (Sarin et al. 2005). We therefore assessed TRT-1-dependent changes in sRNAs throughout the genome, we counted the number of 22G reads mapping uniquely antisense to protein-coding genes and found upregulation of sRNAs at 720 loci and downregulation at 456 loci in *trt-1* animals compared with wild-type controls (Figure S7B). Of the upregulated loci, 465 were not upregulated in *trt-1; dcr-1* animals compared with wildtype, indicating DCR-1 dependence. We did not find significant GO terms in loci with either up or downregulated sRNA production. Given that *trt-1* is expressed at high levels in *C. elegans* germ cells (Meier et al., 2006), we asked whether the loci with differential sRNA production were enriched for germline-enriched genes. We used previously published data on germline expression (Ortiz et al., 2014) and found significant enrichment for germline-enriched genes in both upregulated and downregulated loci (p < 1e-19 and p < 1e-33 respectively, Pearson’s Chi Square test with Bonferroni correction) (Figure S7C). This may reflect an epigenetic response to deficiency for telomerase in germ cells, even though early-generation *trt-1* mutants from which the RNA was prepared had normal levels of fertility. sRNAs mapping to transposons were unaffected by loss of TRT-1, with the exception of the CER-10 LTR retrotransposon, for which sRNAs were three-fold downregulated (Figure S7D).

We found that levels of telomeric sRNAs with 1 to 3 mismatches did not change substantially in small RNAs from *trt-1* single mutants or *trt-1; dcr-1* double mutants, but detected no small RNAs composed of perfect telomere repeats in any genetic background (Figure 5A). We next examined 22G RNAs at subtelomeric regions (defined as 2kb of sequence adjacent to each telomere) and found that these were present in wild-type samples at the left end of chromosome *III*, and that their abundance was increased two-fold in *trt-1* single mutants and their presence almost entirely eliminated in *trt-1; dcr-1* double mutants (Figure 5B, C). Subtelomeric sRNAs were also observed within 1kb of the telomere at the left ends of chromosomes *I* and *IV* in *trt-1* mutants and not in wildtype or *trt-1; dcr-1* double mutants (Figure 5D, E). The right ends of Chromosomes *I* and *III* and the left end of Chromosome *II* all showed at least a two-fold increase in subtelomeric sRNAs in *trt-1* and/or *trt-1; dcr-1* double mutants (Figure 5B).

We observed a significant bias for subtelomeric sRNAs whose sequences correspond to the C-rich telomere strand (antisense to TERRA) for the left ends of Chromosomes *I* and *III* (p < 1e-13 and p < 1e-43 respectively, binomial test with Bonferroni correction) and for the right arm of Chromosome *I* (p < 1e-172, binomial test with Bonferroni correction) while the nucleotide sequences of subtelomeric sRNAs at the left arm of Chromosome *IV* and the right arm of Chromosome *III* showed a bias for subtelomeric TERRA, suggesting that they were created from ARRET (p < 1e-11, p = 0.006 respectively, binomial test with Bonferroni correction) (Figure 5F). These observations suggest that telomere dysfunction triggers the upregulation of small RNA biogenesis at multiple subtelomeres through both DCR-1-dependent and –independent mechanisms. Small RNA biogenesis might be induced at a subset of subtelomeres when telomerase is dysfunctional because 1) shortened telomeres inherited from *trt-1* mutant gametes that created *trt-1* single mutants might trigger sRNA biogenesis, or 2) DNA damage that occurs stochastically at specific subtelomeres elicits sRNA biogenesis. Subtelomeric sRNAs that correspond to the G– or C-rich telomeric strand at specific subtelomeres could be related to the presence of transcripts that can be targeted for sRNA biogenesis at these telomeres.

### Telomere mutations do not explain lack of perfect telomeric small RNAs

The above results are consistent with the possibility that loss of telomerase results in telomere damage and RDRP-mediated biogenesis of small RNA antisense to subtelomeric TERRA at several telomeres. Given this, we were puzzled at the absence of perfect telomeric sRNAs in our *trt-1* mutant sRNA data set. We asked if telomere mutations might explain the absence of perfect telomeric sRNAs by examining telomeres of the VC2010 *C. elegans* strain that is derived from Bristol N2. The VC2010 genome was created by a combination of Oxford Nanopore and PacBio Hifi long read sequencing technologies (Yoshimura et al. 2019). We observed that VC2010 telomeres were covered with mutations, especially at their 5’ ends where TERRA might be transcribed (Figure 6). We next performed PacBio sequencing of our N2 wild type strain, which we obtained from the *Caenorhabditis* Genetics Center in 2001, and which is the genetic background that our *trt-1(ok410)* deletion was back-crossed against prior to performing the experiments reported in this paper. We thawed our N2 wild type strain and passaged two sibling plates in parallel for 60 generations prior to preparing high molecular weight genomic DNA, which was subjected to PacBio Hifi long read sequencing at 35– to 50-fold coverage. We examined telomeres of the genome of the N2 wild type strain SX3254 from Cambridge, England, created by PacBio Hifi sequencing, which confirmed that most N2 wild type strains possess few telomere mutations (Figure 6). De novo genome assemblies were produced using Hifiasm, yielding 35-40 contigs per genome that captured all six chromosome pairs, including 12 telomere ends in each assembly for each N2 genome. We observed a few mutations at 5’ and 3’ ends of some telomeres of both of our N2 genomes, but telomeres were largely free of mutations at their 5’ ends (Figure 6). Moreover, no 3’ telomere mutations were shared between any N2 wild type strain, even our N2 telomeres that were only separated by 50 generations. We conclude that telomerase often acts at *C. elegans* telomeres. In addition, dense telomere mutations were present at the 5’ ends of telomeres of the VC2010 N2 wild type strain but not in our N2 wild type strain, so telomere mutations cannot explain the lack of perfect telomeric small RNAs in our small RNA data sets.

**Figure 6:**
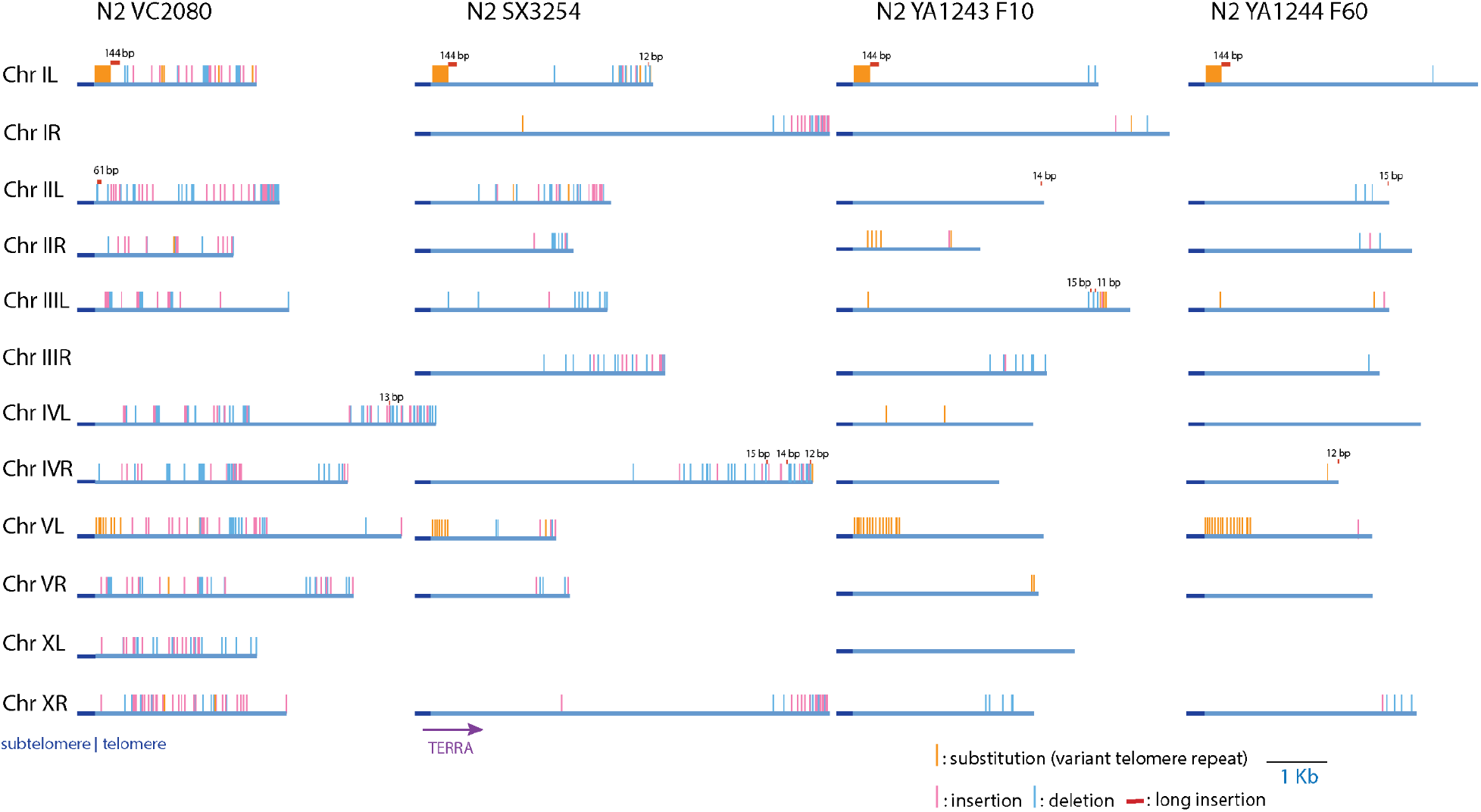
Distribution of telomere mutations in the N2 wild type *C. elegans* strains VC2010, SX3254, Ahmed lab N2 F10, Ahmed lab N2 F60. SNPs and 1-3 nucleotide indel mutations are shown, as well as insertions of >11 nucleotides (red bars). Telomeres were missing from assembled genomic scaffolds for three strains.

### Telomeric sRNAs and loss of EXO-1 repress sterility in the absence of telomerase

TERRA transcription in *S. pombe* initiates in the subtelomere but rarely extends into the telomere (Moravec et al. 2016), so TERRA might serve as a template for RDRP-mediated biogenesis of subtelomeric sRNAs but might rarely yield perfect telomeric sRNAs. It might be difficult to simultaneously target the 11 unique subtelomeres of the *C. elegans* genome with exogenous subtelomeric sRNAs in an effort to show that this class of small RNAs promotes telomere stability in the absence of telomerase. Therefore, we decided to test the hypothesis that perfect telomere repeat small RNAs can promote telomere stability in the absence of telomerase, even though we failed to detect these rare sRNAs in our own sequencing experiments (Figure 5A). We asked if introduction of exogenous telomeric sRNAs could rescue the rapid sterility of sRNA-deficient *trt-1* mutants using a viable *dcr-1* allele, *mg375*, which is deficient for endogenous sRNA production but is hypersensitive to exogenous RNAi when fed *E. coli* expressing dsRNA (Welker et al. 2010). *trt-1; dcr-1(mg375)* double mutant strains that were passaged on bacteria expressing telomeric dsRNA remained fertile significantly longer than those expressing dsRNA from an empty vector control (Figure 7A). In contrast, exogenous telomeric dsRNA did not affect the onset of sterility for *trt-1* single mutant controls. Moreover, *trt-1; dcr-1(mg375)* strains fed telomeric dsRNA had markedly reduced levels of DNA damage signaling at F6 in comparison to *trt-1; dcr-1(mg375)* strains grown on control RNAi or on regular OP50 bacteria that do not express dsRNA (Figure 1F, 7C).

**Figure 7:**
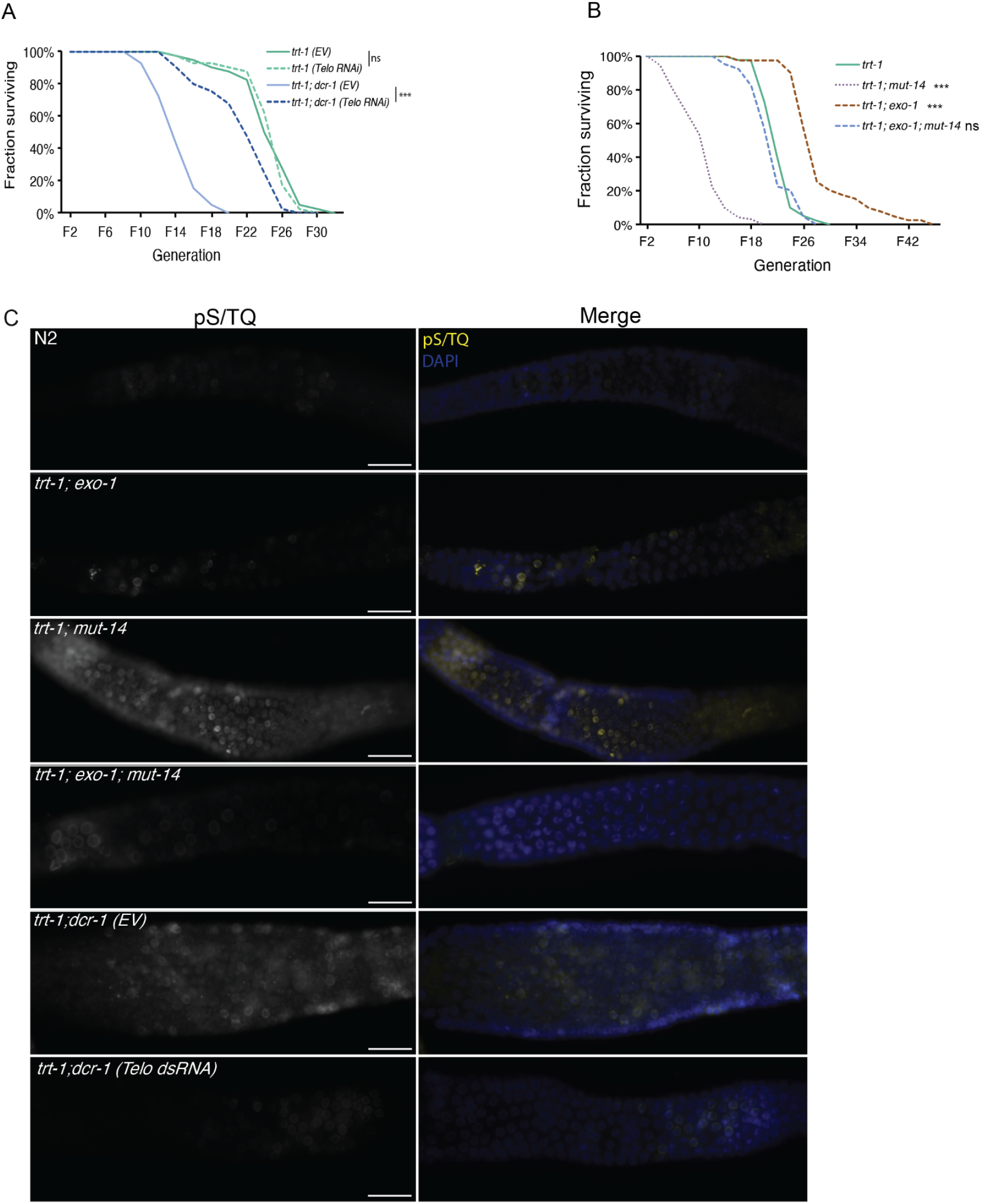
Fast senescence in sRNA-deficient *trt-1* is rescued by exogenous telomeric dsRNA or loss of Exonuclease-1. (A) Survival of *trt-1* or *trt-1; dcr-1(mg375)* on either empty vector control (EV) or bacteria expressing telomeric double-stranded RNA (telo dsRNA). Log rank test relative to EV, ***P < 0.001. n=40 lines per genotype. (B) Loss of *exo-1* extends survival of *trt-1; mut-14* lines. n=40 lines per genotype. (C) Telomeric dsRNA and loss of *exo-1* suppress accumulation of phospho-(Ser/Thr) in sRNAdeficient *trt-1* by F6. Scale bar=20um.

In budding yeast, artificially-induced TERRA overexpression promotes telomere shortening *in cis* by forming RNA:DNA hybrids that stall telomeric replication forks in a manner that stimulates telomere resection by the exonuclease Exo1 (Maicher et al. 2012; Pfeiffer and Lingner 2012). Loss of EXO1 in telomerase-negative mice has been shown to extend lifespan and improve organ maintenance by repressing telomeric ssDNA formation and DNA damage response (Schaetzlein et al. 2007), suggesting a conserved role for EXO1 in the metazoan response to dysfunctional telomeres. We therefore hypothesized that loss of EXO-1 might rescue the accelerated sterility phenotype in small RNA-deficient telomerase mutants. *trt-1; exo-1; mut-14* triple mutant lines survived significantly longer than *trt-1; mut-14* double mutants (Figure 7B) and displayed reduced levels of DNA damage signaling and moderate levels of TERRA at generation F6 (Figures 7C, S7). Moreover, *trt-1; exo-1* double mutant controls remained fertile even longer than *trt-1* single mutants (Figure 7B), indicating that EXO-1-mediated telomere resection promotes telomere-shortening-induced senescence in the absence of telomerase, both at normal telomeres and at telomeres lacking sRNA-mediated genomic silencing. While *trt-1; exo-1* double mutant controls remained fertile longer than *trt-1* single mutants, this was not true for *trt-1* single mutants grown on exogenous telomeric dsRNA (Figure 7A, B). Thus, *exo-1* has at least two roles at telomeres, one that promotes sterility when telomerase is absent and a second that promotes sterility when telomerase and sRNA silencing is absent.

## Discussion

Loss of telomerase and small RNA or heterochromatin factors caused high levels of DNA damage signaling in *C. elegans* germ cells, but only resulted in telomere fusions after ∼10 generations, each generation representing 8-10 cell divisions. Therefore, the accelerated sterility phenotype of small RNA-deficient *trt-1* mutants is not a consequence of a defect in acute telomere uncapping, but rather from accelerated telomere erosion. Stochastic forms of telomere damage that promote telomere erosion include impassable replication blocks caused by stable G-quadruplex structures formed by single-stranded G-rich telomeric DNA during telomere replication (Bryan 2020; Johnson and Brad Johnson 2020; Tsai et al. 2022). In the absence of small RNA-mediated heterochromatin deposition in telomeres of *trt-1* mutants, we observed increased variance in telomere size that could be due to an error-prone form of DNA repair. Our data imply that expression and subsequent small RNA-mediated silencing of TERRA in telomerase mutants promotes favorable repair of telomere damage (Figure 8B).

**Figure 8:**
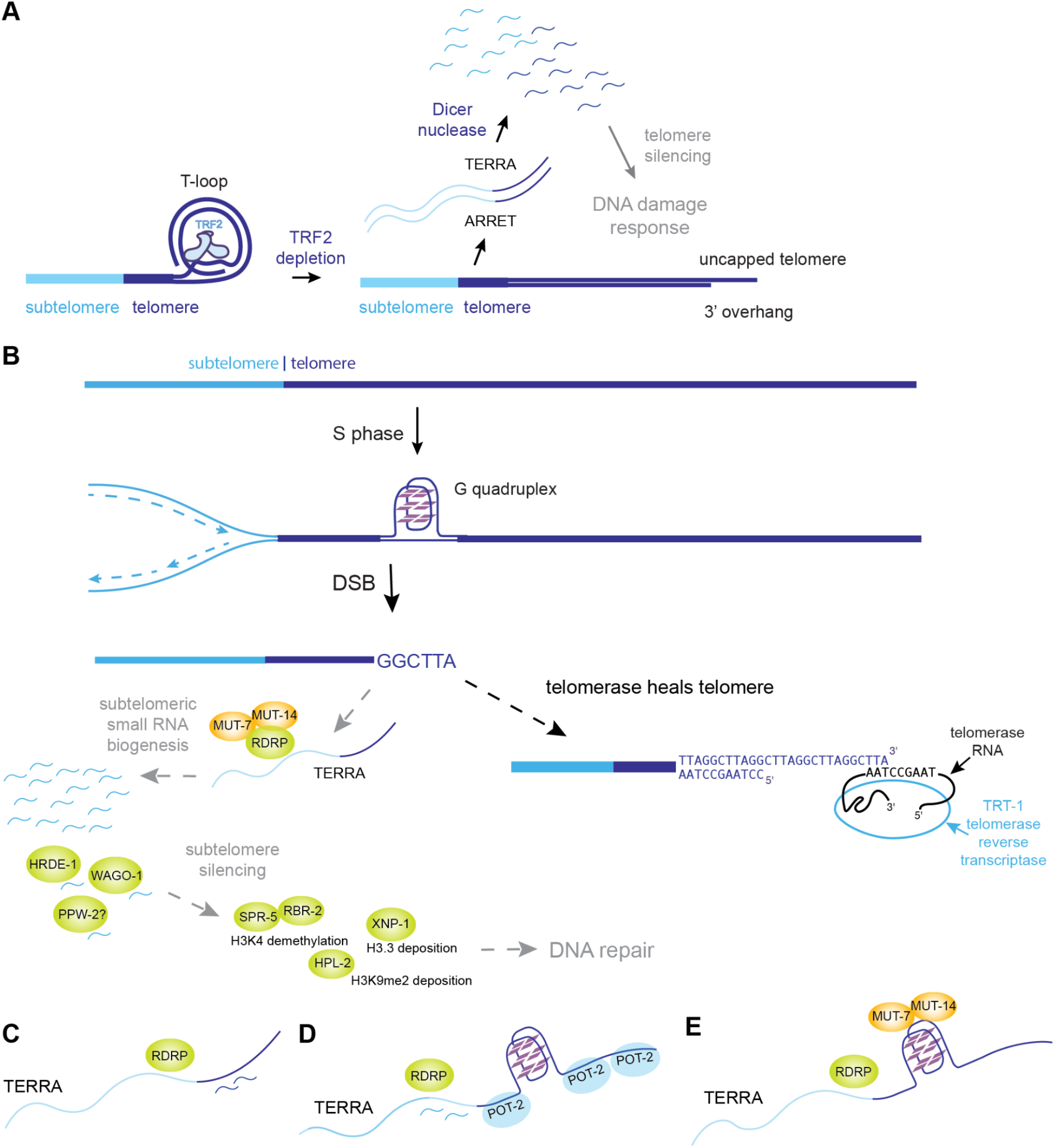
Model for TERRA, small RNAs and telomerase. A) Model from human cells (Rossiello et al. 2017), where inhibition of TRF2 leads to telomere uncapping, TERRA and ARRET expression, and Dicer processing of telomeric dsRNA to create telomeric small RNAs that promote DNA damage response. B) In contrast, *C. elegans trt-1* telomerase mutants produce sub-telomeric small RNAs that interact with Argonaute proteins to promote deposition of H3K9me2 and H3K4 demethylation in order to repress TERRA expression and promote DNA repair. C) Telomeric small RNAs with mismatches could recruit RDRP to TERRA. D) Subtelomeric small RNAs could recruit RDRP to TERRA, if telomere sequence of TERRA is occluded by POT-2 or G-quadruplexes. E) Telomere sequence of TERRA could form a G-quadruplex structure that is sufficient to recruit TERRA to Mutator bodies.

Deficiency for Mutator body proteins MUT-7 or MUT-14 results in loss of RNA dependent RNA polymerase-mediated small RNA biogenesis and lack of effector 22G small RNAs that promote telomere stability in the absence of telomerase. MUT-7 is a homolog of the exonuclease domain of human WRN1 Werner’s syndrome protein whose loss results in early-onset aging of multiple tissues (Ketting et al. 1999). WRN1 deficiency is accompanied by accelerated telomere erosion and senescence phenotypes that can be rescued by telomerase expression (Shimamoto et al. 2015). While WRN1 and its homologs have been intensively studied for their roles in DNA recombination, WRN1 also safeguards the stability of heterochromatin marks (Zhang et al. 2015), which our study suggests promote telomere stability in the absence of telomerase.

Loss of the Mutator protein MUT-16 results in depletion of both subtelomeric small RNAs and telomeric small RNAs with mismatches. We also found that telomeric and subtelomeric small RNAs that associate with HRDE-1 and WAGO-1 Argonaute proteins become depleted in *hrde-1* and *wago-1* mutants. Although WAGO-1 localizes to perinuclear P-granules (germ granules), HRDE-1 localizes to the nucleus where it promotes transcriptional gene silencing (Buckley et al. 2012). Both WAGO-1 and HRDE-1 act in germ cells to silence genes, transposons, and pseudogenes (Gu et al. 2009), but they do not physically interact with one another or possess common interacting proteins (Shirayama et al. 2014; Akay et al. 2017).

We did not observe changes in the levels of telomeric small RNAs in telomerase mutants. We hypothesized that frequent mutations apparent in telomeres of the N2 wild type *C. elegans* strain VC2080 might explain the very low levels of perfect telomeric sRNAs. However, we observed very few telomere mutations for the SX3254 N2 wild type strain of the Miska lab in Cambridge, England, and very few telomere mutations for the Ahmed lab N2 wild type strain at 10 or 60 generations of growth (Figure 6). The lack of perfect telomeric small RNAs in our own data sets, in the presence or absence of telomerase, is consistent with very few perfect telomeric small RNAs in ∼70 sRNA datasets previously queried (Frenk et al. 2019) or in new data sets studied in this paper (Uebel et al. 2018; Seroussi et al. 2023; Knittel et al. 2025). We conclude that small RNAs are almost never created from telomere repeats of TERRA in *C. elegans*.

We previously found that a majority of telomeric small RNAs with mismatches that can be mapped to unique positions in the genome map to one ITS tract, ITS_60. This ITS tract is annotated to create sense and antisense ∼200 nucleotide non-coding RNAs, Y51F10.12 and Y51F10.13, which could anneal to create telomeric dsRNA that is processed by Dicer to promote biogenesis of telomeric small RNAs (Frenk et al. 2019). However, neither RNAi knockdown of *dcr-1* nor *dcr-1(mg375)* mutation nor loss of the DCR-1-interacting protein RDE-4 that binds dsRNA altered the level of telomeric small RNAs with mismatches. This pool of telomeric small RNAs may be maintained, independent of Dicer activity, when ITS tract RNA is expressed during development and targeted by RNA-dependent RNA polymerase. Small RNA pools that target other segments of heterochromatin, such as ancient transposons, may be similarly maintained (Zhang et al. 2011).

Our results are consistent with a role for TERRA expression in the DNA damage response at telomeres discussed by several studies. First, in mammals TERRA expression can occur in response to a distinct form of telomeric damage that occurs in the presence of telomerase – acutely deprotected, non-replicating telomeres – and is accompanied by production of telomeric sRNAs that promote DNA damage response factor recruitment (Rossiello et al. 2017). Second, a study in *S. cerevisiae* showed that increased TERRA expression and persistent telomeric R-loops in telomerase mutants can repress senescence by promoting recombination, indicating that increased levels of TERRA are coupled to induction of a telomeric DNA damage response (Graf et al. 2017; Zeinoun et al. 2023). Third, elevated levels of TERRA have been shown to contribute to senescence in *S. cerevisiae* (Wanat et al. 2018).

Telomere uncapping in somatic mammalian cells results in TERRA expression accompanied by Dicer-dependent telomeric small RNA production (Figure 8A) (Rossiello et al. 2017). This model is consistent with observations of small RNAs that are created at sites of DSB formation in internal segments of the genome (Francia et al. 2012; Wei et al. 2012; Sharma and Misteli 2013). We observed upregulation of small RNAs at some subtelomeres in telomerase mutants, and in some cases this increase was DCR-1-dependent (Figure 5). The variable small RNA biogenesis patterns observed at distinct subtelomeres of telomerase mutants may reflect the epigenetic response to stochastic telomeric DNA damage events. However, we did not observe any perfect telomeric small RNAs in N2 wild type, *trt-1* single mutant or *trt-1; dcr-1(mg375)* double mutant strains (Figure 5). *hrde-1* and *mut-16* small RNA mutant data sets that were wild type for telomerase revealed loss of subtelomeric small RNAs and of telomeric small RNAs with mismatches. However, telomeric or subtelomeric small RNA levels were not reduced for data sets of *rde-4* or *rde-1* single mutants or when *dcr-1* was knocked by RNAi (Table 2). We cannot explain why telomeric and subtelomeric small RNA levels depend strongly on HRDE-1, WAGO-1 and PPW-2 Argonaute proteins and on MUT-16 when telomerase is wild type, under conditions where DCR-1, RDE-4 and RDE-1 do not affect these small RNAs.

Our study reports that subtelomeric small RNAs, generally antisense to TERRA, are induced by *C. elegans trt-1* telomerase mutants. It is possible that RDRP-mediated biogenesis of subtelomeric small RNAs occurs when telomeric small RNAs with mismatches, created from RNA transcribed from ITS tracts, interact with telomere repeats of TERRA at damaged telomeres of *trt-1* mutants (Figure 8C). Alternatively, telomere repeats of TERRA could resist interactions with telomeric small RNAs if telomere repeat RNA is coated with single-stranded telomere binding proteins like POT-1, POT-2, POT-3 or MRT-1, which contain OB-folds that may bind single-stranded telomeric DNA (Raices et al. 2008; Meier et al. 2009; Shtessel et al. 2013; Yu et al. 2023), or if telomere repeat RNA forms G-quadruplex structures that resist interactions with telomeric small RNAs (Figure 8D). This would allow subtelomeric small RNAs to interact with TERRA and recruit RDRP. RNA interference in *C. elegans* requires the RDE-3 ribonucleotidyltransferase, which adds poly(UG) tails to the 3’ ends of mRNAs that are targeted for silencing. Poly(UG) tails form an unusual left-handed G-quadruplex structure that recruits RNA-dependent RNA polymerase to synthesize small RNAs. Therefore, another explanation for lack of perfect telomeric small RNAs is that (TTAGGC)_n_ telomere repeats in *C. elegans* TERRA form an unusual G-quadruplex structure that is sufficient to recruit RDRP to TERRA (Figure 8E) (Marquevielle et al. 2022).

We conclude that subtelomeric small RNAs are likely to interact with TERRA at sites of telomere damage, where they may promote H3K9me2 deposition and facilitate repair of telomere damage (Figure 8B). In this context, telomere repeats of TERRA strongly resist small RNA biogenesis (Figure 8C-E).

## Data Availability

### Accession numbers

RNA-seq data reported in this study are available under the accession number GEO: GSE111800. Genome assemblies of N2 wild type strains YA1243 and YA1244 are available under NCBI accession numbers PRJNA1359394 and PRJNA1403021, respectively.

## Acknowledgements

We thank T. Montgomery for tinyRNA pipeline mentoring, J. Ahringer for heterochromatin comments, R. Dowen for technical advice with qRT-PCR, K. Billmyre for advice on immunofluorescence, and members of the S.A. laboratory for critical review of the manuscript. Some strains were provided by the *Caenorhabditis* Genetics Center, which is funded by National Institutes of Health Office of Research Infrastructure Programs (P40 OD010440). L.L., A.B., S.F., C.L., M.G., E.L.-S. and S.A. were supported by National Institute of Health grants R01 GM135470 and ES035777.

## Competing Interests

The authors declare no competing interests.

## Supplemental Information

7 Supplemental Figures

6 Supplemental Tables

**Figure S1.**
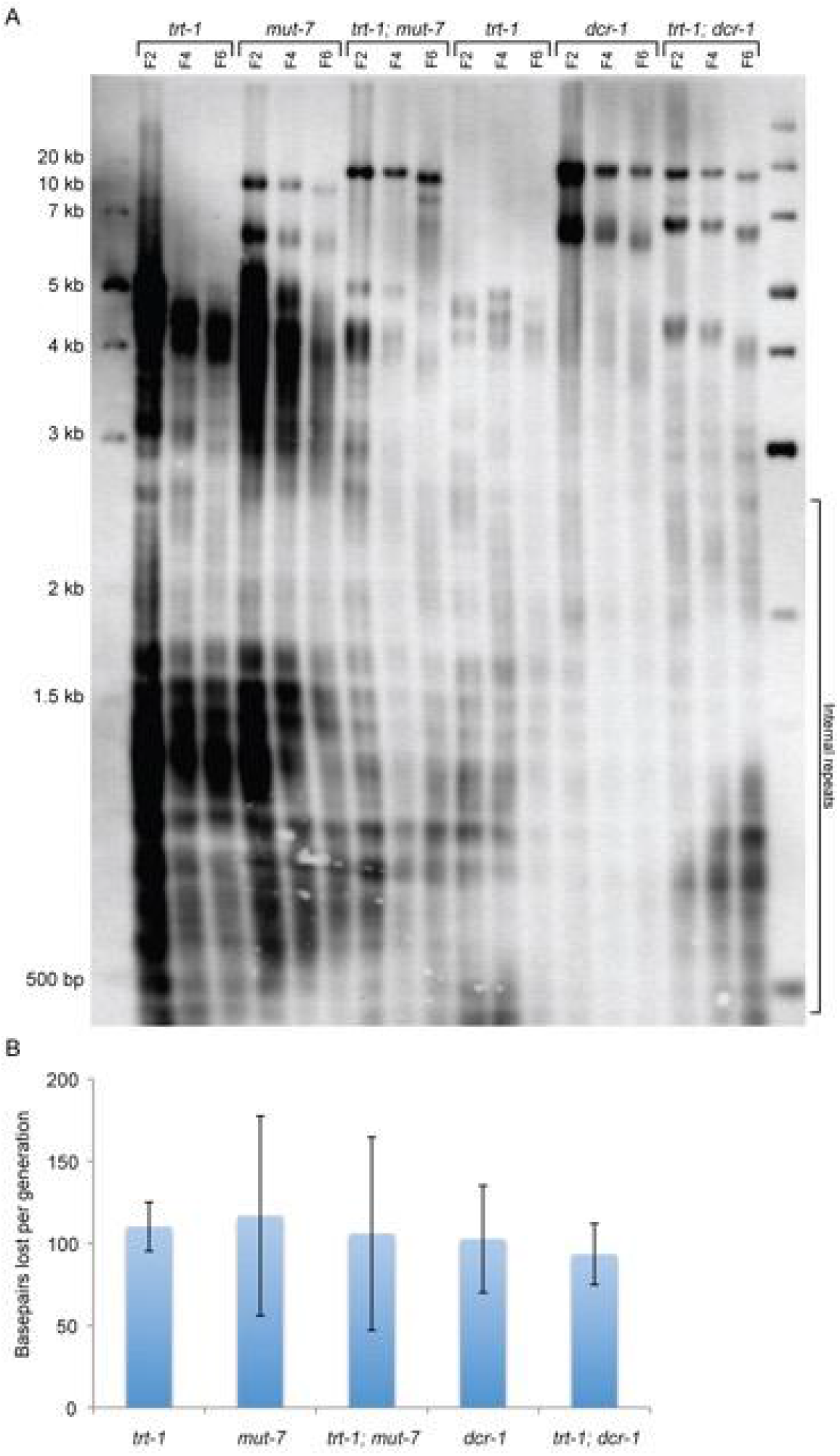
Related to Figure 1. Terminal Restriction Fragment (TFR) analysis of single-mutant *trt-1*, *mut-7*, and *dcr-1* and double-mutant *trt-1; mut-7* and *trt-1; dcr-1* strains using a telomeric probe. (A) Southern blot of samples generated by passaging small groups of worms for multiple generations. Two independent *trt-1* lines were used. Long telomeres observed in strains derived from *mut-7* or *dcr-1* mutant background could be due to rare stochastic telomere elongation events in these backgrounds. (B) Quantification of telomere shortening rate for each genotype shown in panel A. No significant difference was found between any genotypes by Mann Whitney U test.

**Figure S2.**
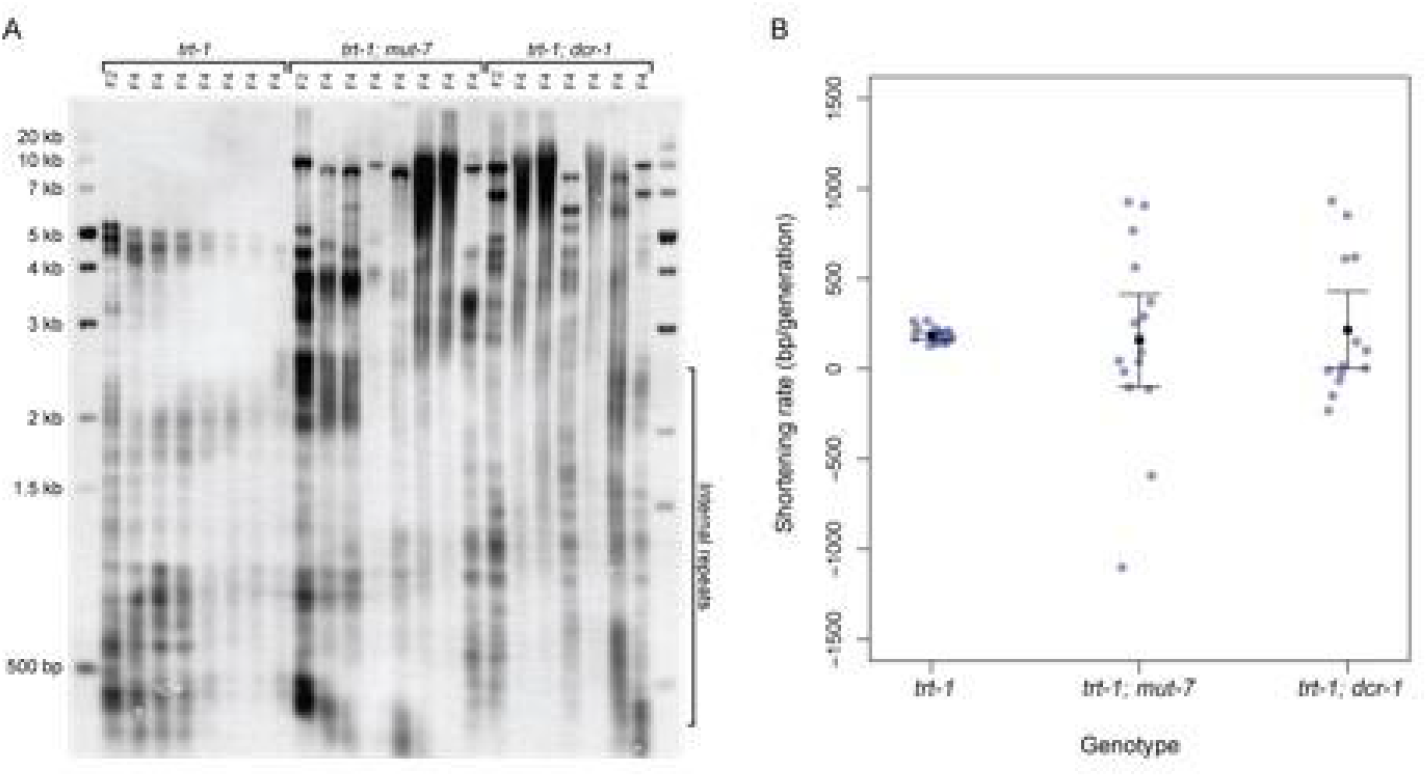
Related to Figure 1. TRF analysis of RNAi-deficient telomerase mutant strains using telomeric probe. (A) A single parental F2 plate was used to generate multiple F4 plates derived from single worms. F4 *trt-1* single mutant lanes show consistent bands corresponding to slight shortening of parental bands. In contrast, F4 lanes from *trt-1; mut-7* and *trt-1; dcr-1* show highly variable bands that in some cases cannot be traced to an obvious parental band. (B) Quantification of telomere shortening rate based on pairing bands appearing at F2 with the closest bands appearing at F4. Highly smeared lanes were excluded. Each point represents one pair of bands. Comparing *trt-1* and *trt-1; mut-7* or *trt-1; dcr-1*, mean shortening rate was not significantly different (p= 0.84 and 0.77, Welch’s t-test) but variance was significantly higher for both double-mutants (p=7.2e-15 and 1.5e-12, F-test). Variance was not significantly different between the double-mutants (p=0.32).

**Figure S3.**
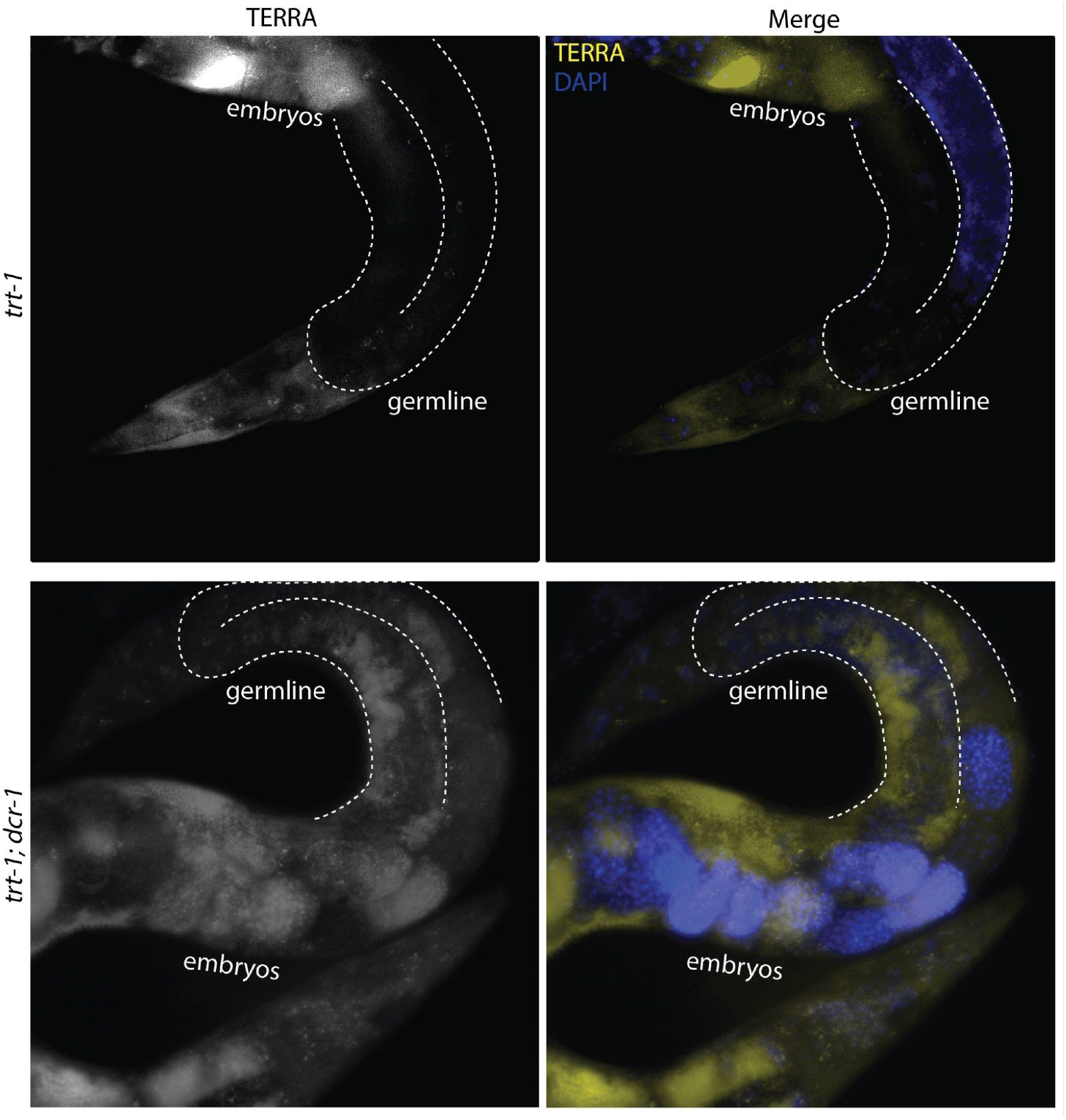
Related to Figure 3. TERRA expression (yellow) in *trt-1* and *trt-1; dcr1(mg375)* 1-day-old adults detected with a Cy5-labelled telomeric probe.

**Figure S4.**
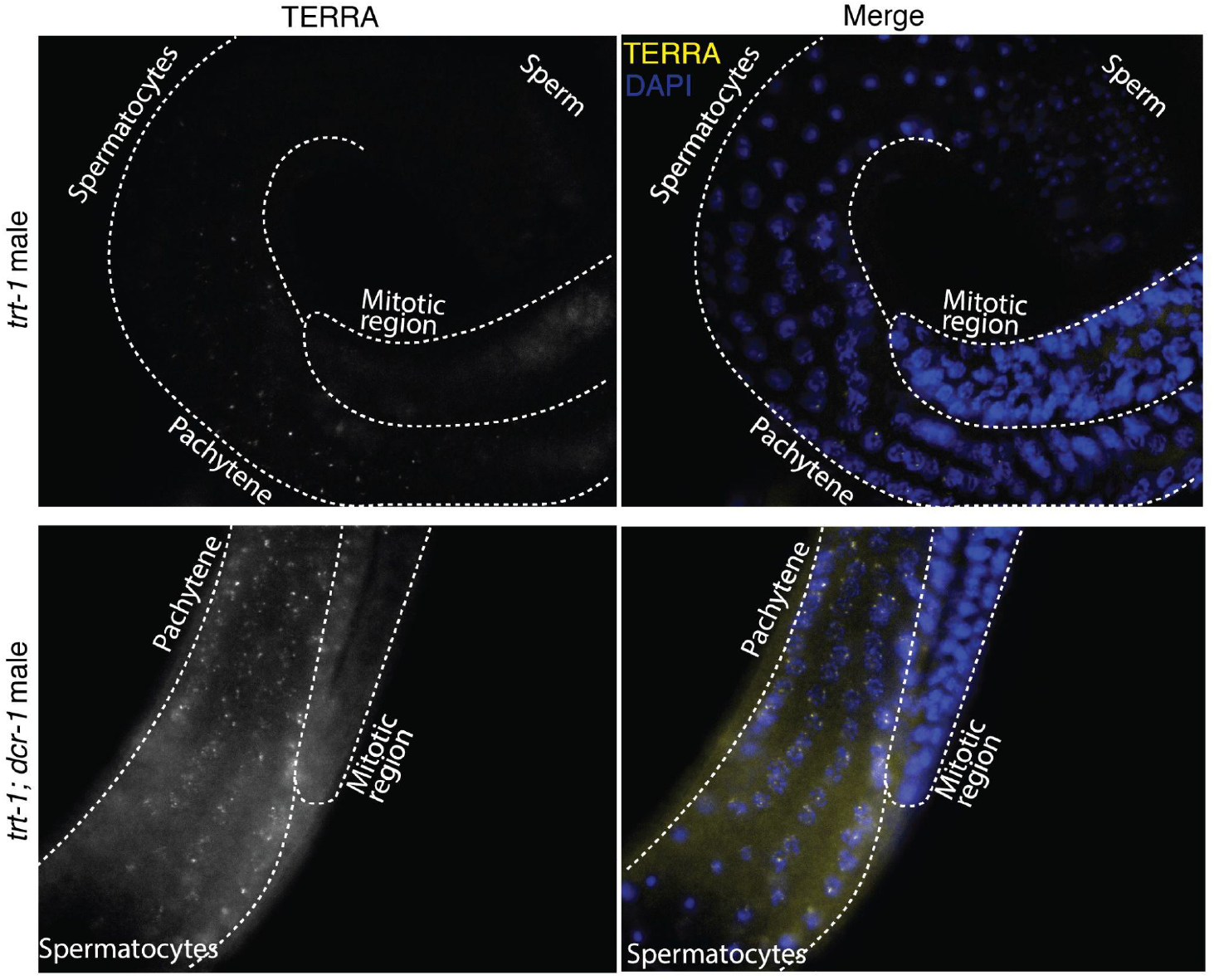
Related to Figure 3. RNA FISH of TERRA expression in *trt-1* and *trt-1; dcr1* males. Note TERRA foci in the Pachytene region in both *trt-1* and *trt-1; dcr-1*, as well as lack of expression in the mitotic region and spermatocytes.

**Figure S5.**
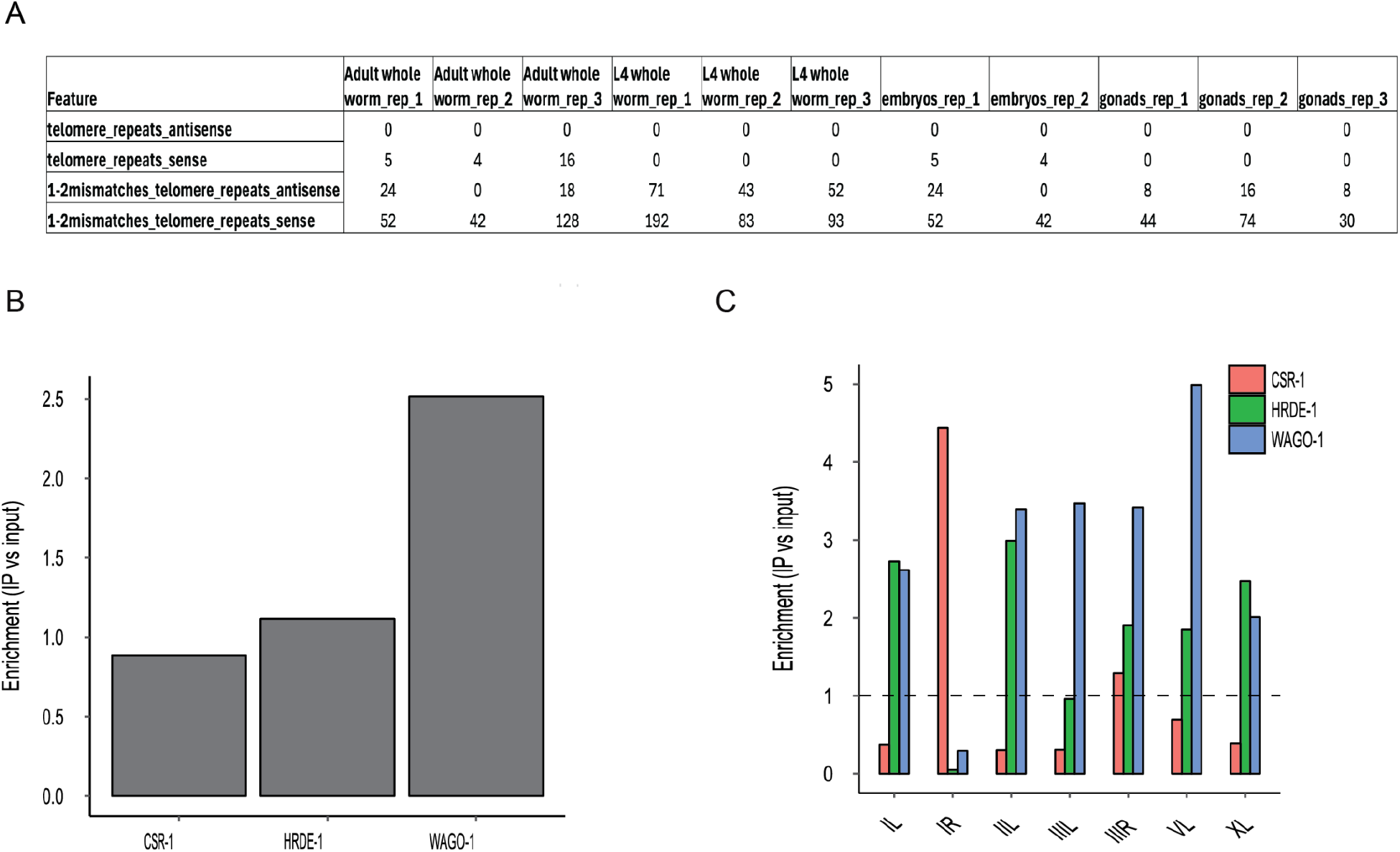
Related to Figure 3. Telomeric and Subtelomeric small RNA analysis on CSR-1, HRDE-1, and WAGO-1 IP small RNA datasets. (A) Perfect matches to telomeric repeats are rare. Read counts of small RNAs matching seven telomeric repeats with 0 or up to 2 mismatches. Results were generated using tinyRNA analysis pipeline. (B) Enrichment (IP/input) of reads mapping to perfect telomere sequence with up to three mismatches in published immunoprecipitation datasets for three different Argonaute proteins. (C) Enrichment for subtelomeric sRNAs.

**Figure S6.**
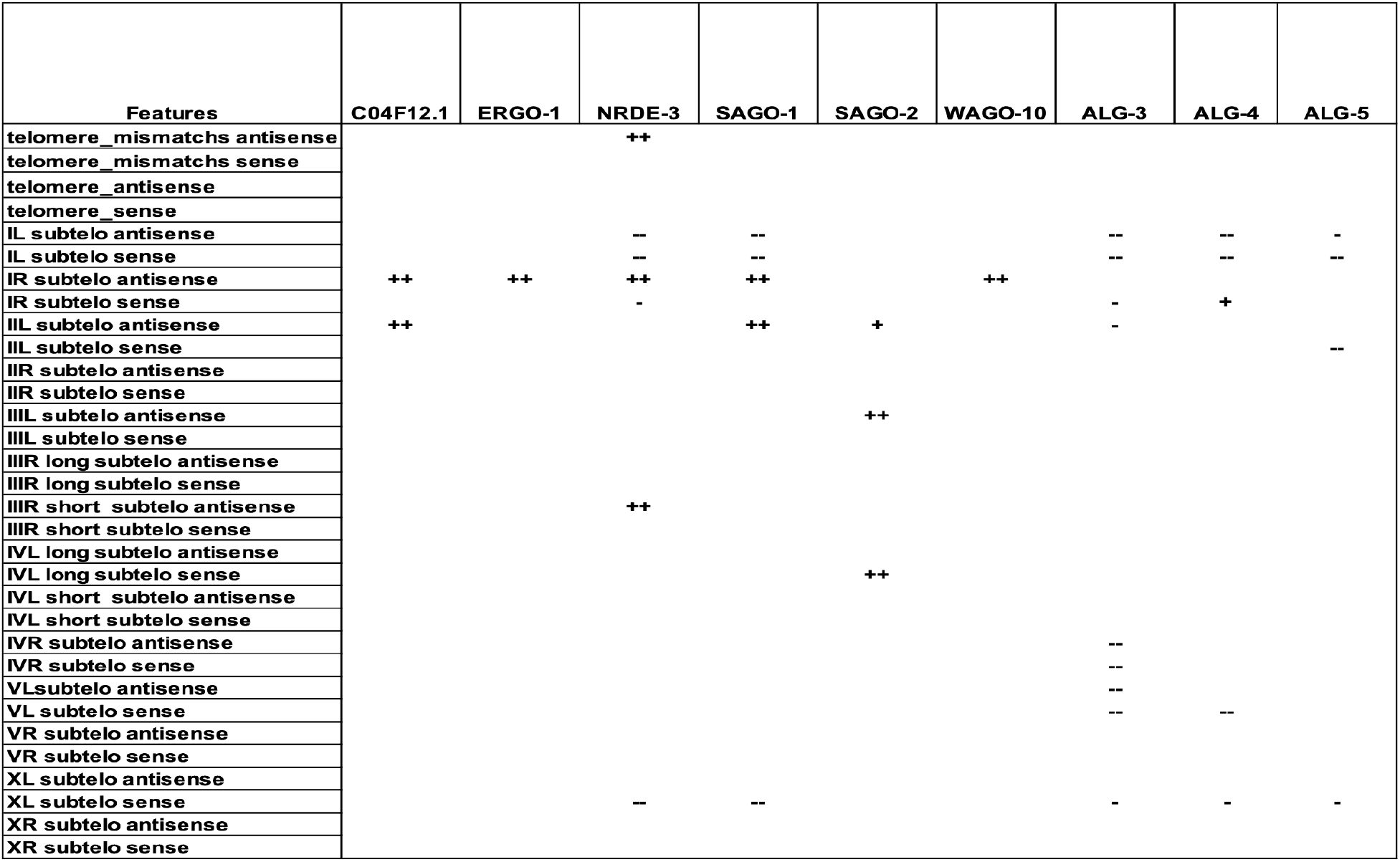
Related to Figure 4. A subset of Argonaute IP small RNA sequencing datasets showed no telomeric or subtelomeric enrichment. Symbols indicate differential expression using DESeq2, “+”/“-” denote log2 fold change > 1 with p-adjusted < 0.05, and “++”/“--” denote log2 fold change > 2 with p-adjusted < 0.05.

**Figure S7.**
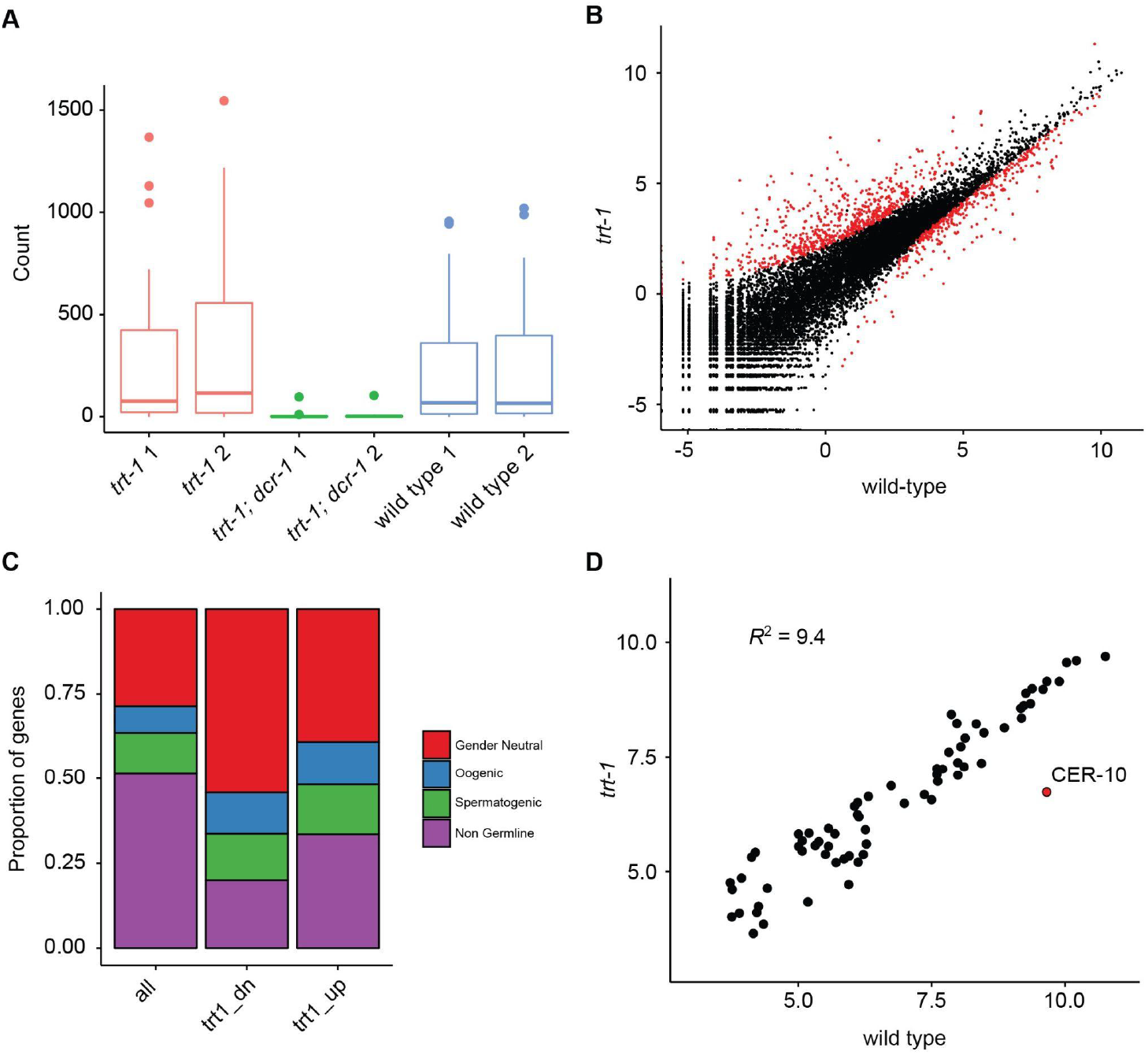
Related to Figure 5. Analysis of 22G small RNA species in telomerase and Dicer mutants. (A) Number of reads mapping to DCR-1 targets identified by Gent *et al*. (Gent et al., 2010). (B) log2 normalized mean read counts at every *C. elegans* gene for *trt-1* and wildtype. Genes identified as significantly differentially expressed by DESeq2 are shown in red. (C) Proportion of all genes or genes downregulated (trt1_dn) or upregulated (trt1_up) belonging to expression categories defined by Ortiz *et al*. (see reference 54 in main text). (D) log2 normalized mean read counts at transposon consensus sequences for *trt-1* and wildtype.

**Supplemental Table 1.** Related to Figure 1. Statistical analysis of ALT survival in double mutant strains compared to single mutant *trt-1* control. The Z score represents how many standard deviations from the number of ALT survivors in double mutants is from the *trt-1* single mutant control mean survival rate. The p-value represents the probability that the number of survivors via ALT is not significantly different from the control *trt-1* single mutant strain.

|  | Lines tested | Number ALT survivors | Z score | p-value (two tailed) |
|---|---|---|---|---|
| *trt-1* | 40 | 6 |  |  |
| *trt-1; mut-7(pk204)* | 40 | 4 | 0.68 | 0.50 |
| *trt-1; mut-14(pk738)* | 40 | 4 | 0.68 | 0.50 |
| *trt-1 xnp-1(ok1823)* | 40 | 6 | 0 | 1 |
| *trt-1 xnp-1(tm678)* | 40 | 4 | 0.68 | 0.50 |
| *trt-1; dcr-1 (mg375)* | 40 | 5 | 0.32 | 0.75 |

**Supplemental Table 2.** Related to Figure 1. Log rank analysis of survival curves of double or triple mutants compared to single mutant *trt-1* controls. n=40 independent lines for each genotype. Strains were passaged by transferring 6 larval L1s weekly and scored as sterile when plates contained no new L1 larvae. The p-value represents the probability that there is no difference in the probability of sterility at any time point between the double or triple mutant strain and the *trt-1* single mutant strain.

| Strain | Log Rank test (Z=) | p-value |
|---|---|---|
| *trt-1; mut-7(pk204)* | 9.79 | <0.00001 |
| *trt-1; mut-14(pk738)* | 9.96 | <0.00001 |
| *trt-1; dcr-1(mg375)* | 9.37 | <0.00001 |
| *trt-1; rde-1* | 8.23 | <0.00001 |
| *trt-1; rde-4* | 9.21 | <0.00001 |
| *trt-1; rrf-3* | 8.4 | <0.00001 |
| *trt-1; rrf-2* | 1.7 | 0.089 |
| *trt-1; exo-1* | 7.38 | <0.00001 |
| *trt-1 xnp-1(ok1823)* | 4.3 | <0.00001 |
| *trt-1 xnp-1(tm678)* | 4.23 | <0.0001 |
| *trt-1 xnp-1(tm678); mut-7* | 9.21 | <0.00001 |
| *trt-1; sid-1* | 6.52 | <0.00001 |
| *trt-1; rbr-2(tm1231)* | 8.64 | <0.00001 |
| *trt-1; spr-5 (by101)* | 8.89 | <0.00001 |

**Supplemental Table 3.**
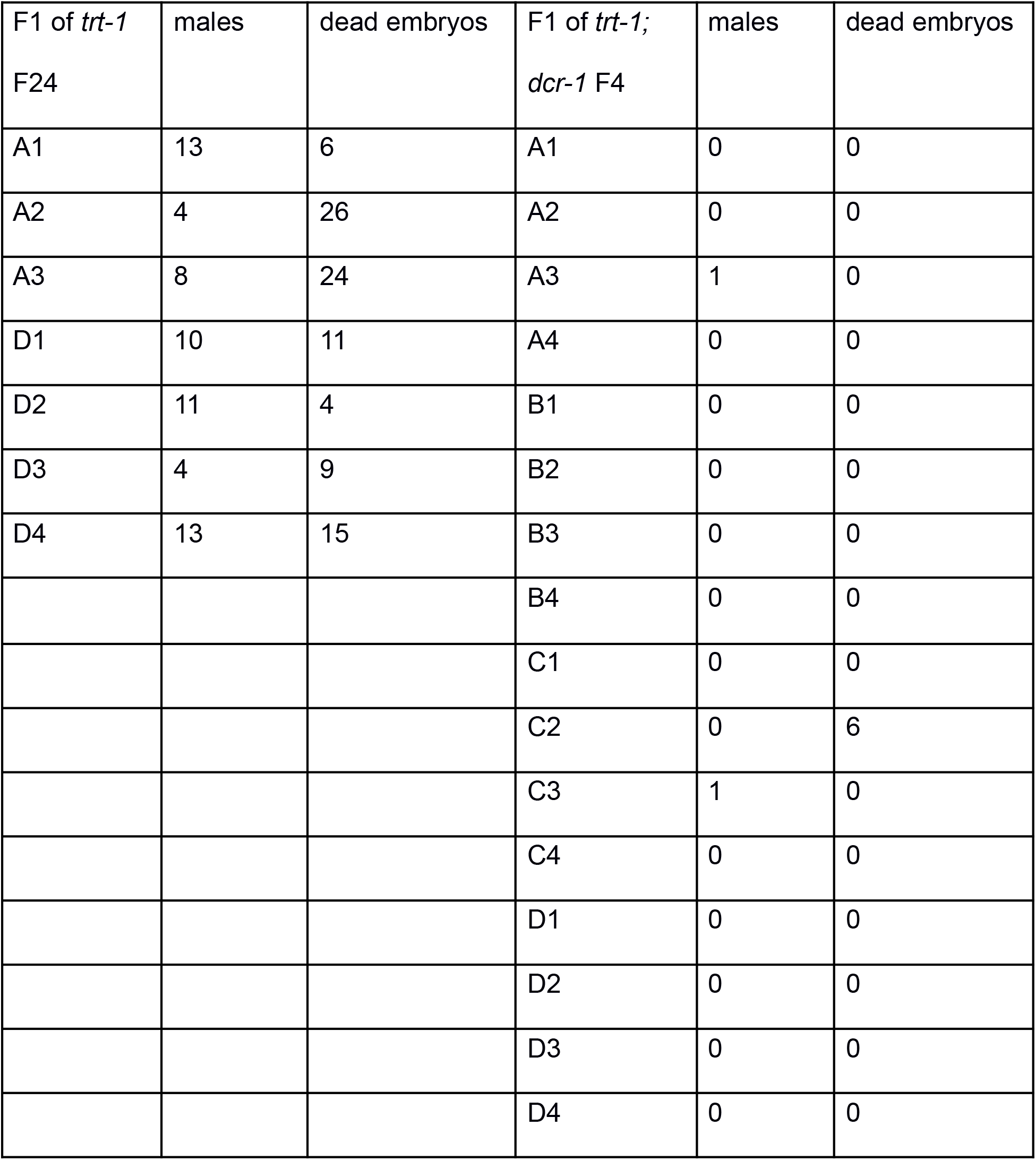
Related to Figure 1. Counts of total F2 males and dead embryos scored for F1 progeny from crosses of wild-type males with either late generation F16 *trt-1* mutant controls (n=2 independently grown strains, three F1 scored for strain A and four F1 for strain D) or early generation F4 *trt-1; dcr-1* double mutant (n=4 strains, 4 F1 each scored for strains A, B, C, and D) hermaphrodites. If a chromosome fusion is present in any of the F1 animals, then it will cause weak nondisjunction of the fused chromosomes, which leads to F2 dead embryos if an autosome is fused or to F2 males if the X chromosome is fused.

**Supplemental Table 4.** Related to Table 1. Differential expression analysis of telomeric and subtelomeric small RNAs on N2 datasets.

**Supplemental Table 5.** Related to Table 1. Differential expression analysis of telomeric and subtelomeric small RNAs on Argonaute Immunoprecipitation small RNA sequencing datasets.

**Supplemental Table 6.** Related to Table 2. Differential expression analysis of telomeric and subtelomeric small RNAs on datasets with small RNA pathway disruption.

## Notes

### Competing Interest Statement

The authors have declared no competing interest.

### Summary of Updates

We have extensively rewritten the manuscript in response to Reviewer comments, including substantial shortening and refocusing of the Discussion and rewording of the abstract.

